# Sex influences gliovascular unit assembly and function in the developing mouse brain

**DOI:** 10.64898/2026.04.13.718096

**Authors:** Lucie Lemale, Myriam Abioui-Mourgues, Lydie Naulé, Rodrigo Alvear-Perez, Marina Rubio, Denis Vivien, Mathilde Becmeur-Lefebvre, Tristan Hourcade, Anne-Cécile Boulay, Sakina Mhaouty-Kodja, Martine Cohen-Salmon, Barbara Delaunay-Piednoir

## Abstract

The gliovascular unit (GVU, a specialized interface between the brain and the vascular system) assembles and matures after birth and establishes essential homeostatic functions, including blood–brain barrier integrity, metabolic exchanges, fluid drainage, neurovascular coupling, and immune surveillance. Here, we compared the postnatal maturation of the cortical GVU in male vs. female mice. On P15, males exhibited more angiogenesis, a higher vessel density, and a higher level of aquaporin 4 expression in perivascular astrocyte processes. Females exhibited a higher density of perivascular macrophages and fibroblasts on P15 and P30 respectively, earlier development of the arterial vascular network, and a greater capacity to recruit cerebral blood flow. Transcriptomic profiling during the P5–P120 period revealed sex-specific developmental trajectories within the GVU, with the most prominent differences observed on P5. These dimorphisms were regulated at least in part by oestrogenic signalling. Taken as a whole, our results highlight pronounced sex-dependent differences in GVU assembly and maturation that might influence brain physiology and vulnerability to neurodevelopmental disorders.

## Introduction

Recent evidence indicates that sex differences influence neurodevelopmental trajectories and contribute to susceptibility to and the prevalence and clinical presentation of many neurological and neurodevelopmental disorders. For example, autism spectrum disorder, attention-deficit/hyperactivity disorder, and several cognitive disorders are more prevalent in males, whereas migraine, depression, and multiple sclerosis are more prevalent in females ^1–7^. Sex differences have also been reported in neurovascular and neurodegenerative diseases, including stroke and Alzheimer’s disease^8–10^. Although genetic, hormonal and environmental factors probably contribute to these disparities, the underlying biological mechanisms have not been fully characterized. Importantly, the gliovascular unit (GVU, a multicellular interface which includes blood vessels (ranging from arteries and veins to capillaries) and astrocytes) regulates blood–brain barrier (BBB) integrity, cerebral blood flow (CBF), neurovascular coupling, immune surveillance, and brain homeostasis – all of which are implicated in the pathophysiology of neurological disorders^11–13^. Sex-specific differences in the maturation of GVU components during critical developmental periods might influence disease susceptibility later in life. However, despite the growing recognition of sex differences in brain development and disease, the extent to which GVU maturation differs between males and females has not been systematically investigated. This knowledge gap has been further exacerbated by the historical underrepresentation of females in biomedical research^14^.

Given the GVU’s central role in brain development, homeostasis, and neurological disease, we sought to determine whether its postnatal maturation is influenced by sex. To this end, we characterized the different cellular compartments of the murine and human GVUs and analysed the GVU transcriptomes in males and females during postnatal development and in adulthood. Our results reveal sex-specific developmental trajectories within the GVU and provide new insights into the biological basis of sex differences in brain development.

## Results

### On postnatal day (P)15, angiogenesis is more active in male mice than in female mice

The cerebral vasculature is immature at birth and so undergoes substantial postnatal remodelling and growth. A recent study in C57BL/6J mice identified three sequential phases of postnatal cerebral vascular development: an expansion phase (extending until P7), a regional specialization phase (up to P21), and a refinement phase (from P21 onward, characterized by vascular network stabilization and maturation)^15^. Here, we sought to determine whether these developmental processes exhibit sex-specific differences. We first compared males and females with regard to the endothelial architecture during postnatal development by immunolabeling the endothelial protein CD31 (also known as Pecam1) on coronal brain sections (**Fig. 1a**). We focused our analysis of the somatosensory cortex on three timepoints: P5 (during vascular expansion), P15 (when angiogenesis and vascular remodelling are both active), and P30 (in the maturation phase) ^15^. We quantified the length of the vessels, the number of segments, the angle of the branches, and segment straightness. (**Fig. 1b-g**). A huge increase in vessel density was observed between P5 and P15, which corresponds to the postnatal angiogenic phase described above ^16,17^ (**Fig. 1c-e**). On P15, the vascular network was denser in males than in females (**Fig. 1c-e**). The vessel straightness was similar in males and females at all stages (**Fig. 1g**). No differences in any of the variables were detected on P30 (**Fig. 1c-g**).

**Figure 1:**
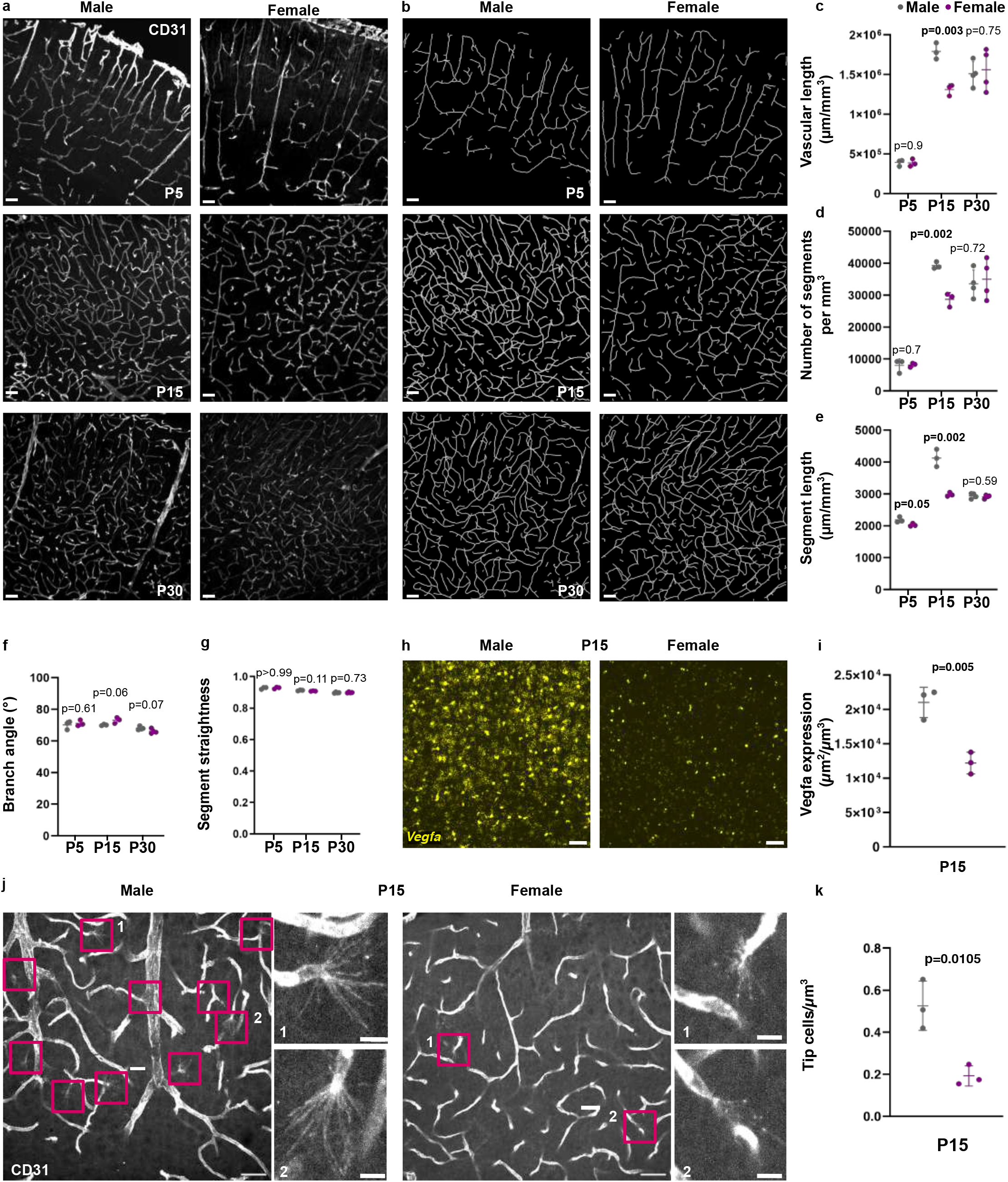
Postnatal angiogenesis is more active in male mice than in female mice on P15. **a**. Representative spinning-disk microscopy images of the somatosensory cortical endothelial network immunolabeled for CD31 (in grey) on P5, P15 and P30 in males and females. Scale bars: 50 µm. **b.** A 3D reconstruction of the filaments corresponding to vessels. Scale bars: 50 µm. **c-g.** Quantification of the vascular network in the somatosensory cortex. Length **(c),** number of segments **(d);** segment length (**e**); branch angle (**f**); segment straightness (**g**). n_P5 male_ =3, n_P5 female_ = 3, n_P15 male_ = 3, n_P15 female_ = 3, n_P30 male_ = 4, n_P30 female_ = 4 (2-4 images per mouse). The data are presented as the mean ± SD and were analysed using an unpaired t-test for (**c-f**) and a Mann-Whitney test for (**g**). **h.** Representative spinning-disk images of *Vegfa* mRNA detected by FISH (in yellow) in somatosensory cortex sections on P15. Scale bars: 50 µm. **i.** Quantification of *Vegfa* mRNA normalised against the field of view volume. n= 3 for each sex (4 images per mouse). The data are presented as the mean ± SD and were analysed using an unpaired t-test. **j**. Representative images of vascular network immunolabeled for CD31 (in grey) on P15, with the presence of tip cells (indicated by red squares). Enlarged views of tip cells 1 and 2 are shown on the right. Scale bars: 50 µm. **k.** Quantification of tip cells normalised against the field of view volume. n= 3 for each sex (4 images per mouse in the somatosensory cortex). The raw data are given in **Source Data Table S5.**

We next investigated the mechanisms underlying the denser vascular networks observed in males on P15. Brain endothelial development and architecture is regulated by the pericytes recruited to capillary endothelial cells during embryogenesis ^18^. We used CD13 immunolabelling to analyse pericyte vascular coverage of the capillary network on P5, P15, and P30 **(Fig. S1a-d).** Vessels were counterstained with isolectin-B4 (IB4). The coverage of CD13 immunolabeling on IB4 was identical in males and in females at all stages of development **(Fig. S1e)**. We next assessed the level of mRNA expression of *Vegfa*; this gene codes for a potent angiogenic factor, the level of which is correlated with vessel expansion^15^. We detected *Vegfa* mRNA by fluorescent *in situ* hybridization (FISH) on cortical sections on P15 (**Fig. 1h**). The *Vegfa* expression level was markedly higher in males than in females (**Fig. 1i**). Interestingly, the somas with *Vegfa* dots were mainly positive for the astrocyte-specific transcription factor Sox9 in males (**Fig. S1f**) but not in females (**Fig. S1g**). Lastly, on P15, we observed a higher number of CD31^+^ endothelial tip cells sprouting from vessels in the somatosensory cortex (indicative of ongoing angiogenesis) in males than in females (**Fig. 1j, k**).

These results show that angiogenesis in the somatosensory cortex remains more active at P15 in males in females and thus results in a transiently denser vascular network.

### On P15, astrocyte density and perivascular levels of aquaporin 4 (Aqp4) are higher in males than in females

Astrocytes are generated by the final differentiation of radial glial cells around the time of birth. In rodents, astrocytes proliferate and colonize the brain during the first postnatal week ^19^. The cells then extend, ramify, and take on a “bushy” shape in contact with synapses and blood vessels. In the somatosensory cortex, perivascular astrocyte processes (PvAPs) cover 25% of the capillaries on P0, 50% on P5, 75% on P7, 95% on P10, and close to 100% on P15 ^20–22^. Here, we analysed astrocyte development in males and females. We first analysed astrocyte density on P5, P15, P30, and P60 in males and females by using Sox9 immunolabelling of cortical sections to detect astrocyte somas in the somatosensory cortex (**Fig. 2a**). Due to discrepancies that have been described in the literature ^23,24^, we counted Sox9^+^ astrocyte somas with *cellpose* (https://github.com/orion-cirb/Orion_Macros.git), a deep-learning-based segmentation method (**Fig. 2b**). As observed previously ^19,25^, the number of Sox9 somas was higher on P5 than on P15 – indicating strong glial reorganization during this time window. Notably, the number of Sox9^+^ astrocytes was higher in males than females on P15 (**Fig. 2c**).

**Figure 2:**
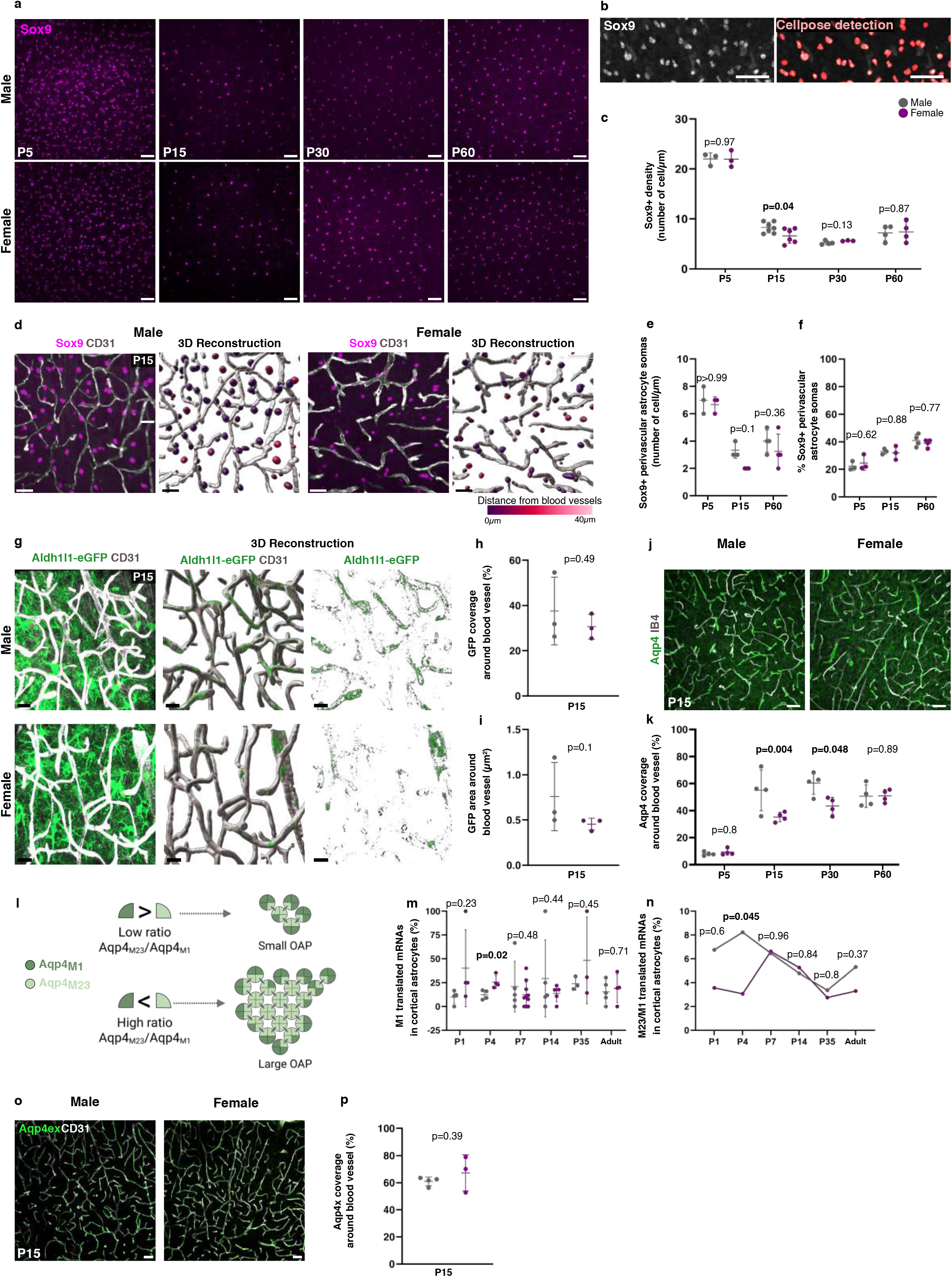
Astrocyte density and perivascular levels of aquaporin 4 (Aqp4) are higher in males on P15. **a.** Representative images of astrocyte somas immunolabeled for Sox9 (in magenta) in male and female somatosensory cortex on P5, P15, and P30. Scale bar: 50 µm. **b. left**, Representative image of Sox9 immunolabelling (in grey) on P5; **right,** detection of Sox9 signals by *cellpose*. Scale bars: 50 µm. **c.** Quantification of Sox9^+^ cells. n_P5 male_ = 3, n_P5 female_ = 3, n_P15 male_ = 7, n_P15 female_ = 6, n_P30 male_ = 4, n_P30 female_ = 3, n_P60 male_ = 4, n_P60 female_ = 4 (4 images per mouse). The data are presented as the mean ± SD and were analysed using an unpaired t-test. **d.** Representative images of astrocyte somas immunolabeled for Sox9 (in magenta) and vessels immunolabeled for CD31 (in grey) in male and female somatosensory cortex on P15 with 3D reconstruction. Scale bar: 30 µm. **e.** Quantification of the density of Sox9^+^ astrocyte somas within 0.8 µm of a blood vessel (perivascular astrocyte somas). **f.** Percentage of perivascular astrocyte somas. n_P5 male_ = 3, n_P5 female_ = 3, n_P15 male_ = 3, n_P15 female_ = 3, n_P60 male_ = 4, n_P60 female_ = 4 (2-4 images per mouse). The data are presented as the mean ± SD and were analysed using an unpaired t-test or a Mann-Whitney test for (**e**) and a Chi² test for (**f**). **g.** Representative images of astrocyte cytoplasmic eGFP immunolabeling in Aldh1l1-eGFP transgenic male and female somatosensory cortex on P15. Vessels are immunolabelled for CD31 (in grey). On the rigth, 3D reconstruction of eGFP perivascular signal. Scale bar: 20 µm. **h.** Quantification of the percentage of perivascular eGFP signal. **i.** Quantification of the area of perivascular eGFP. n_P15 male_ = 3, n_P15 female_ = 3. The data are presented as the mean ± SD and were analysed using an unpaired t-test for (**h**) and a Mann-Whitney test for (**i**). **j.** Representative spinning-disk images of astrocyte perivascular processes immunolabelled for Aqp4 (in green) in male and female somatosensory cortex sections on P15. Vessels are stained with IB4 (in grey). Scale bars: 50 µm. **k.** Quantification of perivascular Aqp4 on P5, P15, P30 and P60. n= 4 for each sex and each stage (4 images per mouse). The data are presented as the mean ± SD and were analysed using a chi^2^ test. **l**. Schematic representation of Aqp4 OAP size dependency on the M1 and M23 Aqp4 isoform composition. **m.** Quantification of the percentage of translated M1 mRNAs in cortical astrocytes and the M23/M1 translated mRNA ratio calculated from Rurak et al. ^24^ **n.** Representative spinning-disk images of astrocyte perivascular processes immunolabelled for Aqp4ex (in green) in male and female somatosensory cortex sections on P15. Vessels are immunolabeled for CD31 (in grey). Scale bars: 50 µm. **o.** Quantification of perivascular Aqp4ex. n_males_= 4 and n_females_= 3 (4 images per mouse). The data are presented as the mean ± SD and were analysed using an unpaired t-test. The raw data are given in **Source Data Table S5.**

We next looked at whether these sex differences in Sox9 immunolabelling were reflected on the gliovascular level. Firstly, we measured the distance between Sox9^+^ soma and brain vessels immunolabelled for CD31 on P5, P15, and P30 (**Fig. 2d**). No significant differences were observed at any stage (**Fig. 2e-f**). Furthermore, astrocyte perivascular coverage in the somatosensory cortex (reflected by the level of cytoplasmic perivascular eGFP on Aldh1l1:eGFP cortical sections) was identical in males and females on P15 (**Fig. 2g-i**).

It was recently reported that neocortical astrocytes achieve a mature phenotype earlier in males than in females ^24^. We sought to determine whether this phenomenon influenced the PvAPs, which display a highly dynamic postnatal molecular maturation process ^26^. Here, we focused on Aqp4, a water channel that is enriched in PvAPs and is crucial for perivascular homeostasis ^27^. On cortical sections, vessels were stained with IB4, and astrocytes were immunolabelled for Aqp4 (**Fig. 2i and Fig. S2**). On P5 and P60, Aqp4 perivascular immunolabeling was essentially identical in males and females. However, on P15 and P30, Aqp4-vascular coverage was higher in males **(Fig. 2j)**. In astrocyte perivascular membranes, Aqp4 tetramers form orthogonal arrays of particles (OAP) composed of Aqp4 M1 and M23 isoforms (produced by alternative splicing ^28^ (**Fig. S3a**)) and Aqp4ex isoforms (M1ex, and M23ex, in which a 29-amino-acid C-terminal extension is generated by translational readthrough) ^29–31^. M23 is about five times more abundant than M1, and this ratio is mandatory for the formation of OAPs ^32,33^. The size of the OAPs is proportional to the M23/M1 ratio (**Fig. 2k**). Aqp4ex is essential for Aqp4’s localization in the astrocyte perivascular membrane ^29,31^. We first analysed the translatome database of neocortical astroglial cells ^24^, in order to quantify the alternative exon junctions corresponding to M1 and M23 in male and female neocortical astrocytes. Interestingly, the proportion of M1 isoforms on P4 was higher in astrocytes from females than in astrocytes from males – suggesting that the M23/M1 ratio was lower (**Fig.2 l-m; Fig. S3b**). The level of Aqp4ex immunodetected on cortical sections on P15 was essentially identical in males and females (**Fig. 2 n, p**).

Taken as a whole, the results show that male mice have more astrocytes than female mice on P15. Although the astrocyte perivascular coverage is ultimately similar in the two sexes, males transiently display more perivascular Aqp4 – perhaps as a result of the developmental expression profile of the M1Aqp4 isoform.

### The recruitment of Lyve 1^+^ perivascular macrophages (PVMs) on P15 and *Col1a1*^+^ perivascular fibroblasts (PVFs) on P30 is more intense in females than in males

Brain PVMs and PVFs are emerging as essential regulators of cerebrovascular development, maturation, and homeostasis; they are thought to coordinate vascular integrity, extracellular matrix remodelling, and neurovascular interactions throughout brain development and into adulthood ^34,35^. PVFs and PVMs emerge from leptomeninges during the second postnatal week and are recruited in the parenchymal perivascular spaces around large, penetrating arterioles and exiting venules, with which they interact closely^36,37^. PVFs express a large variety of basement membrane ECM proteins ^35^ and contribute to the dynamic properties of vascular membranes ^38,39^. PVMs are crucial for the regulation of immunity, cerebrospinal fluid drainage, and CBF ^40,41^. PVM recruitment is known to be influenced by PVFs ^42^. Both PVMs and PVFs are implicated in diseases (including amyotrophic lateral sclerosis ^43^, stroke, and Alzheimer’s disease) and ageing ^44–47^.

In the present study, we quantified the PVF and PVM density in the somatosensory cortex of males and females on P15, P30 and P60 by using (i) FISH to detect *Col1a1 mRNAs* expressed specifically by PVFs, and (ii) immunolabelling to detect the PVM reference markers CD206 and Lyve-1 ^32^ (**Fig. 3a)**. The detection of a very bright *Col1a1* signal around large, penetrating, cortical vessels enabled us to count individual cells **(Fig. 3a)**. We observed more PVFs in females transiently, on P30 (**Fig. 3b**). For PVMs, and although the number of CD206□ cells was similar in males and females (**Fig. 3c**), we observed a higher number of Lyve1□ PVMs in females than in males on P15 (**Fig. 3d**). This difference reflected a higher fraction of blood vessels harbouring Lyve1□ cells (**Fig. 3e**). Moreover, we identified distinct PVM subpopulations ^48,49^ (**Fig. 3f-h)**: CD206^+^ Lyve1^+^ and CD206□ Lyve1□ PVMs had similar profiles in the two sexes (**Fig. 3f-j**), whereas CD206^-^ Lyve1□ PVMs were more abundant in females than in males (**Fig. 3h, k**).

**Figure 3:**
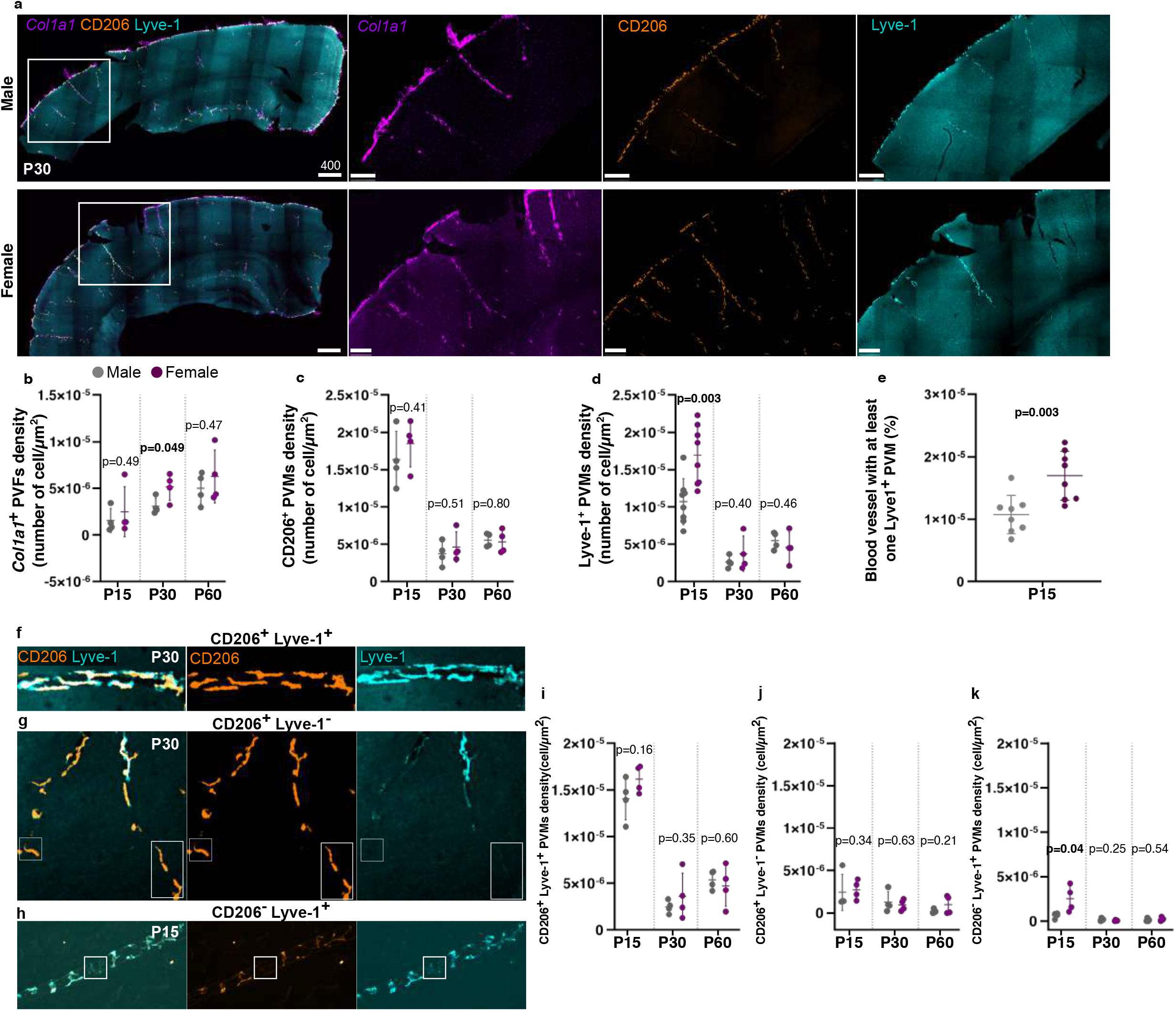
The recruitment of Lyve 1+ PVMs on P15 and *Col1a1* PVFs on P30 is more intense in females. **a**. Representative images showing FISH detection of *Col1a1* in PVFs (orange) and immunolabelling of Lyve-1 (cyan) and CD206 (orange) in PVMs on P30 in somatosensory cortex sections from males and females. Scale bar: 400 µm (left panel) and 100 µm (right panel) **b**. Quantification of *Col1a1*^+^ PVF density on P15, P30 and P60. n= 4 for each sex and each stage. The data are presented as the mean ± SD and were analysed using an unpaired t-test or a Mann-Whitney test. **c**. CD206^+^ PVM density n= 4 for each sex and each stage. **d**. Lyve-1^+^ PVM density. n_P15 male_ = 8, n_P15 female_ = 8, n_P30male_ = 4, n_P30 female_ = 4, n_P60 male_ = 4, n_P60 female_ = 4. **e**. The percentage of blood vessels with at least one PVM in the somatosensory cortex on P15. n_P15male_ = 8, n_P15female_ = 8. The data are presented as the mean ± SD and were analysed using an unpaired t-test. **f**. An enlarged view of a blood vessel with co-immunolabelled CD206^+^Lyve-1^+^ PVM. Scale bar: 50 µm. **g**. An enlarged view of a blood vessel with a CD206^+^ Lyve-1^-^ PVM. Scale bar: 50 µm. **h**. An enlarged view of a blood vessel with a CD206^-^Lyve-1^+^ PVM. Scale bar: 50 µm. **i-k**. Counts of CD206^+^ Lyve-1^+^, CD206^+^ Lyve-1^-^ and CD206^-^ Lyve-1^+^ PVMs. n_male_ = 4, n_female_ = 4. The data are presented as the mean ± SD and were analysed using an unpaired t-test or Mann-Whitney test. The raw data are given in **Source Data Table S5.**

Overall, our results show that males and females differ with regard to the recruitment of PVMs and PVFs in the cortex. Lyve1^+^ PVMs and *Col1a1*+ PVFs are transiently more numerous in females than in males on P15 and P30, respectively.

### On P15, the arteriolar network and the CBF are more developed in females than in males

We had shown previously that VSMCs acquire their contractile properties between P5 and P15 in mice and after birth in humans ^50^. Thus, we assessed the contractile differentiation of VSMCs from P5 to P30 in males and females by using light-sheet microscopy to image immunolabelled smooth muscle actin (SMA) in the somatosensory cortex on cleared brain samples (**Fig. 4a**). As reported previously, the length and complexity of the SMA-positive vascular network increased progressively from P5 to P30 ^50^ **(Fig. 4a-e)**. However, the length and number of SMA^+^ vessels on P15 **(Fig. 4a- d)** and the number of SMA^+^ branches on P30 **(Fig. 4a, d)** were higher in females than in males.

**Figure 4:**
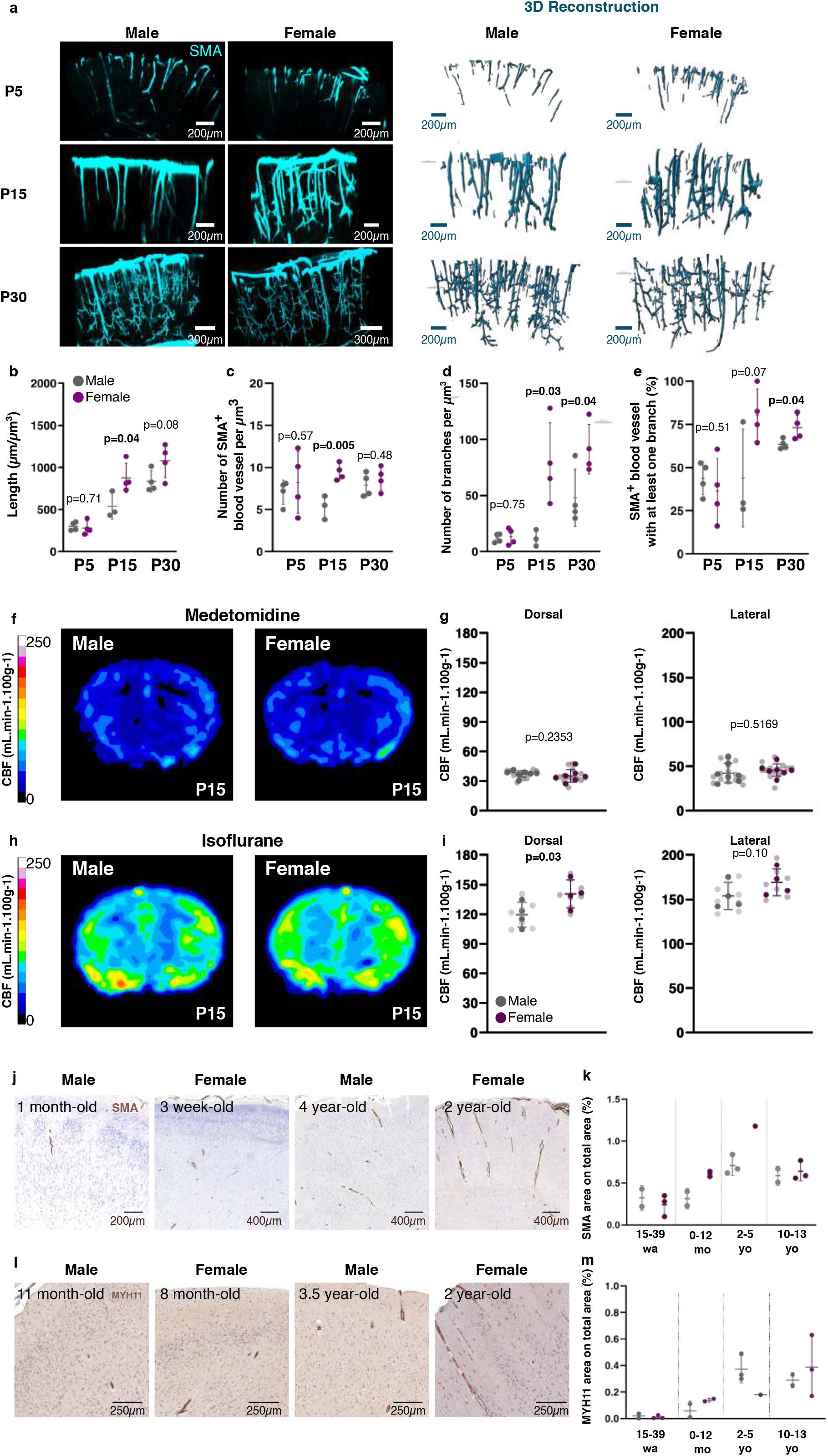
The arteriolar network and the CBF are more developed in females on P15. **a.** Representative light-sheet microscopy images of whole cleared somatosensory cortex immunolabeled for smooth muscle actin (SMA; in cyan) on P5, P15, and P30 in males and females (**left panel**) and their respective 3D reconstructions (**right panel**). **b–e** Quantification of SMA□ blood vessels: total vessel length (**b**); total number of vessels (**c**); number of secondary branches (**d**); number of SMA^+^ vessels with at least one secondary branch (**e**). The data are presented as the mean ± SD and were analysed using an unpaired t-test. n_P5 male_ = 4, n_P5 female_ = 4, n_P15 male_ = 3, n_P15 female_ = 4, n_P30 male_ = 4, n_P30 female_ = 4. **f.** Representative map of *in vivo* arterial spin labelling MRI acquisitions in males and females on P15 under medetomidine anaesthesia (the “awake” condition). **g.** Analysis of CBF in the dorsal and lateral cortex. Dark dots represent the mean value for each mouse, and light dots represent the values for each hemisphere. The data are presented as the mean ± SD and were analysed using an unpaired t-test (dorsal cortex) and unpaired t-test (lateral cortex). n_male_ = 7, n_female_ = 7. **h.** Representative map of *in vivo* arterial spin labelling MRI acquisitions in males and females on P15 under isoflurane anaesthesia (the “vasodilation” condition). **i.** Analysis of CBF in the dorsal and lateral cortex. Dark dots represent the mean value for each mouse, and light dots represent the values for each hemisphere. The data are presented as the mean ± SD and were analysed using a one-tailed unpaired t-test (dorsal cortex) and an unpaired t-test (lateral cortex). n_male_ = 4, n_female_ = 4. **j**. Immunohistochemical analysis of SMA and MYH11 expression in the developing cortex in non-pathologic human males and females. **k**. Quantification of SMA, MYH11 stained surface areas (quoted as the mean ± SD, whenever possible). The low number of samples prevented the application of statistical tests. The raw data are given in **Source Data Table S5**.

Since the development of the SMA^+^ arterial network in the brain parenchymal is correlated with increases in VSMC contractility and CBF ^50^, we next used arterial spin labelling (ASL) magnetic resonance imaging (MRI) to measure CBF in the cortex of male and female mice on P15 ^51^. We did not observe a difference between male and female mice anesthetized with medetomidine, which has only a small effect on vasoreactivity under physiological conditions ^52^ (**Fig. 4f, g**). In contrast, CBF in the dorsal cortex was significantly higher in females than males when the mice were anesthetized with isoflurane, which induces dose-dependent strong pharmacologically-driven cerebral vasodilation ^53^. A non-significant trend was observed in the lateral cortex (**Fig. 4fh, i**).

We had shown previously that in humans, the level of cortical SMA and MYH11 is higher after birth than before birth and continues to increase until the age of 5 years - suggesting that VSMC contractility also matures postnatally in humans ^50^. We re-analysed our previous data by stratifying by sex (**Fig. 4h, j, k; Fig. S4, 5)**. The levels of SMA and MYH11 were the same in the two sexes prenatally but tended to be higher for SMA only in females than in males at 0-2 years of age and at 5 years of age. From the age of 10 years onwards, the levels of both proteins were similar in females and males **(Fig. 4j, k; Fig. S4, 5).**

Overall, our results show that the arterial network is more developed in female mice than in male mice between P15 and P30. This enhanced maturation is associated with a greater capacity to recruit the CBF in response to physiological demands. A similar, sex-specific pattern of postnatal cerebrovascular maturation might occur in humans and might be linked to sex differences in SMA expression.

### Oestradiol modulates development of the endothelium and the arteriolar network

Gonadal hormones induce early brain sexual dimorphism in both rodents and humans. In male mice, two testosterone surges occur before and after birth ^54–56^ and is converted into oestradiol in the brain (**Fig. 5a**). In female mice, the release of oestradiol from the ovaries starts later on P7 ^57,58^ and levels increase progressively to reach a first peak on P12-P14 ^59^. Circulating sex steroids can access the brain through the BBB ^60^. Here, we hypothesized that this early-life hormonal secretion provides a plausible mechanistic framework for the sexualization of GVU development.

**Figure 5:**
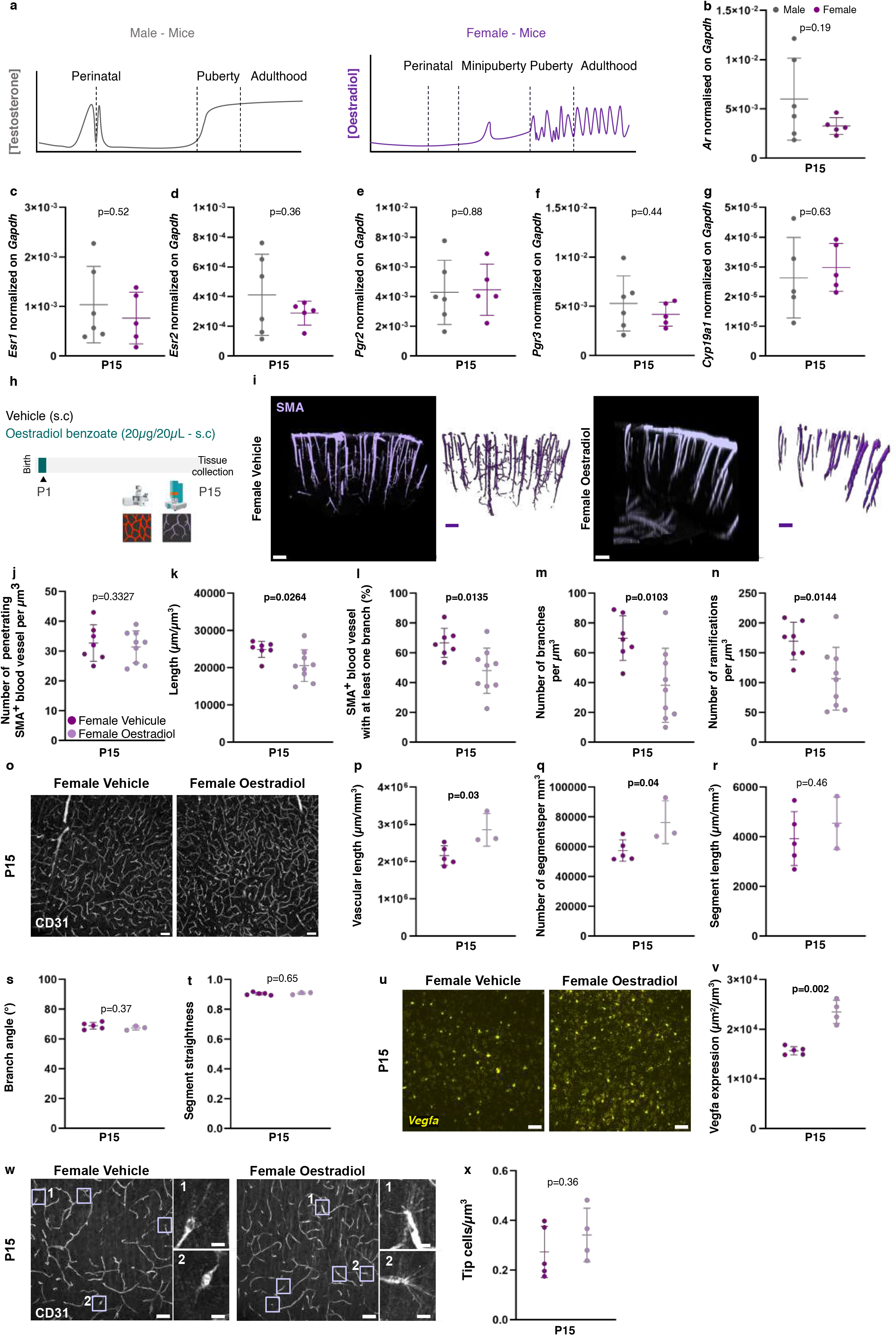
During perinatal development, arteriolar network and vascular growth are regulated by oestrogen signalling. **a.** A schema representing variations in the circulating levels of sex steroid hormones in male (testosterone; left panel) and in females (oestrogen; right panel) during development **b-g.** qRT-PCR analysis of testosterone, oestradiol and progesterone receptors, and *Cyp19a1* encoding aromatase in whole P15 male and female brains. The results were normalised against *Gapdh* expression. n_P15Male_= 5-6, n_P15Female_= 5. The data are presented as the mean ± SD and were analysed using an unpaired t-test. **h.** Female mice received a single subcutaneous injection of oestradiol benzoate (20 µg) on P1. Brains were collected on P15 for light-sheet imaging or spinning disk imaging. **i.** Representative light-sheet microscopy images of whole cleared somatosensory cortex immunolabeled for smooth muscle actin (SMA; in purple) on P15 in vehicle- and oestradiol-benzoate-treated females (**left panel**) and their respective 3D reconstructions (**right panel**). **j-n.** Quantification of SMA□ blood vessels: number of penetrating vessels (**j**); total vessel length (**k**); number of vessels with at least one secondary branch (**i**); number of secondary branches (**m**); number of ramifications (**n**). The data are presented as the mean ± SD and were analysed using an unpaired t-test. n_P15 female vehicle_ = 7, n_P15 female oestradiol benzoate_ = 9 in two independent experiments. **o.** Representative spinning-disk microscopy images of brain sections immunolabeled for CD31 in vehicle- and oestradiol-benzoate-treated females. **p-t.** Quantification of the vascular network: Length (**p**), number of segments (**q**); segment length (**r**); branch angle (**s**); segment straightness (**t**). n_P15 female vehicle_ = 5, n_P15 female oestradiol-benzoate_ = 3. The data are presented as the mean ± SD and were analysed using an unpaired t-test. **u**. Representative spinning-disk images of *Vegfa* mRNA detected by FISH (in yellow) in somatosensory cortex sections in vehicle- and oestradiol-benzoate-treated females. Scale bars: 50 µm. **v.** Quantification of *Vegfa* mRNA normalised against field of view volume. n_P15 female vehicle_ = 5, n_P15 female oestradiol-benzoate_ = 4. The data are presented as the mean ± SD and were analysed using an unpaired t-test. **w.** Representative images of vascular network immunolabeled for CD31 (in grey) in vehicle- and oestradiol-benzoate-treated females, with the presence of tip cells (purple squares). Enlarged views of tip cells 1 and 2 are shown on the right. Scale bars: 50 µm. **x.** Quantification of the number of tip cells normalised against the field of view volume. n_P15 female vehicle_ = 5, n_P15 female oestradiol-benzoate_ = 4. The data are presented as the mean ± SD and were analysed using an unpaired t-test. The raw data are given in **Source Data Table S5**.

We first used quantitative reverse transcription and polymerase chain reaction (RT-qPCR) assays to measure the expression of the androgen receptor (encoded by *Ar*), oestrogen receptors (encoded by *Esr1* and *Esr2*), progesterone receptors (encoded by *PgR1* and *PgR2*), and aromatase (encoded by *Cyp19a1* and which converts testosterone to oestrogen locally)^61^ in the whole brain of males and females on P15, i.e. the timepoint showing the greatest sexual dimorphism. No differences between males and females were observed. (**Fig. 5b-g**).

We next injected newborn females with 20 µg of oestradiol benzoate or (as a control) oil vehicle on P1 and assessed several properties of the GVU on P15 (**Fig. 5h**). We first used light-sheet microscopy of cleared brain samples to assess the SMA-immunolabelled arteriolar network organization in the somatosensory cortex (**Fig. 5i**). Strikingly, and although the number of large, penetrating vessels remained stable over time (**Fig. 5j**), we observed greater falls in the length, ramification and branching of SMA vessels in oestradiol-benzoate-treated females than in control females (**Fig. 5k-n**). We next observed the CD31-immunolabelled vessel network on brain sections (**Fig. 5o**). The vascular density (**Fig. 5p**) and the number of segments (**Fig. 5q**) were both higher in oestradiol-benzoate-treated females than in control females, while the segment length (**Fig. 5r**), branch angle (**Fig. 5s**) and segment straightness (**Fig. 5v**) values were similar. To possibly relate this effect to angiogenesis, we used FISH to assess *Vegfa* expression in oestradiol-benzoate-treated females and control females (**Fig. 5u, v**). The *Vegfa* signal was much stronger in oestradiol-benzoate-treated females than in control females (**Fig. 5u, v**). However, the numbers of CD31^+^ tip cells were similar in the two groups of mice **(Fig. 5 w, x)**. Lastly, oestradiol benzoate treatment did not have a significant effect on Sox9^+^ astrocyte density (**Fig. S6 a, b**) or Aqp4 perivascular immunolabeling (**Fig. S6 c, d**).

These results demonstrate that the injection of oestradiol at birth in females masculinizes the postnatal development of the arterial and vascular systems.

### The GVU transcriptome has different developmental trajectories in males and females

Following our anatomic and functional characterization of the developing GVU, we next sought to investigate its molecular properties. Specifically, we looked at whether GVU development displayed sex-specific transcriptomic signatures. We previously reported that the mechanical purification of brain microvessels (MVs) preserves their cellular content (i.e. endothelial cells, VSMCs, pericytes, PVMs, PVFs, and attached PvAPs) at all stages ^50,62^. These preparations also contain associated neuronal, microglial, and possibly oligodendrocyte processes and their associated mRNAs ^50^. Here, MVs were extracted from whole cortices, and purified mRNAs were sequenced and compared in males vs. females on P5, P15 and P120 **(Fig. 5)**. To confirm that our RNA sequencing (RNA-Seq) libraries were enriched in vascular mRNAs, each gene’s expression pattern was annotated on the basis of a previous, adult, single-cell transcriptome analysis ^63,64^ (**Table S1**; **Fig. S7**). In all libraries, endothelial genes were predominant (**Fig. S7a**). VSMC transcripts and arterial endothelial transcripts were more strongly represented in P120 libraries, which reflects the late postnatal maturation of the arterial system^50^ (**Fig. S7b**). Interestingly, P5 MVs from females had a unique profile, with a lower percentage of endothelial genes and a higher percentage of pericyte genes (**Fig. S7a**).

Genes expressed differentially in males vs. females were defined as those with a log2 fold change (FC) ≤ -1 or log2FC ≥ 1 and an adjusted p-value or false discovery rate ≤ 0.05 (**Table S1**). On P120, only 20 genes were expressed differentially in males vs. females (**Fig. 6a; Table S1**). Nine differentially expressed genes (DEGs) were enriched in males, including *Lcn2* (encoding lipocalin 2, which is expressed in some ECs and is involved in inflammatory processes; FC: 11.24)^65^ and *Ptgs2* (encoding prostaglandin-endoperoxide synthase 2/cyclooxygenase 2, which is expressed mainly by certain PVFs and is involved in the regulation of CBF; FC: 4.00) ^66^. Eleven DEGs were enriched in females, including the *Angptl4* gene encoding angiopoietin-like 4 (a secreted protein that binds to the ECM and regulates angiogenesis; FC: 2.17) ^67^. On P15, no sex differences were observed (**Fig. 6a; Table S1**). In contrast to the results for P15 and P120, 335 male vs. female DEGs were detected on P5. Of these, 306 were upregulated in males (**Fig. 6a; Table S1**). The Reactome pathway analysis on P5 showed that the most strongly regulated pathways were related to vascular functions, such as the basal lamina composition (*Elastic fiber formation*, *Collagen biosynthesis and modifying enzymes*, *Collagen formation* and *ECM organization*) and transport across the BBB (*solute carrier (SLC) mediated transport, Transport of inorganic cations and anions,* and *Transport of small molecules*) (**Fig. 6b**). With regard to *ECM organization*, the genes most strongly enriched in males were *Col8a1* and *Col8a2* (FCs: 10.01 and 23.85, respectively), which are mainly expressed in adult PVFs (**Fig. 6a; Table S1**). *Utrn* (encoding utrophin, a member of a large protein complex linking actin to the membrane and basal lamina and which is involved in smooth muscle contractility; FC: 9.08) was greatly enriched in females over males ^68^ **(Table S1**). With regard to SLCs, the PVF-specific gene *Slc13a4* (encoding a sodium sulphate cotransporter; FC: 26.19) was highly upregulated in males, *Slc1a2* (encoding the glutamate transporter 1; FC: 3.52) was enriched in females, and *Slc39a12* (encoding the zinc transporter protein ZIP12; FC: 4.41) was enriched in males (**Fig. 6a; Table S1**). The last two genes are expressed only in astrocytes and contribute respectively to the perisynaptic uptakes of glutamate and zinc required for optimal neurotransmission ^69,70^.

**Figure 6:**
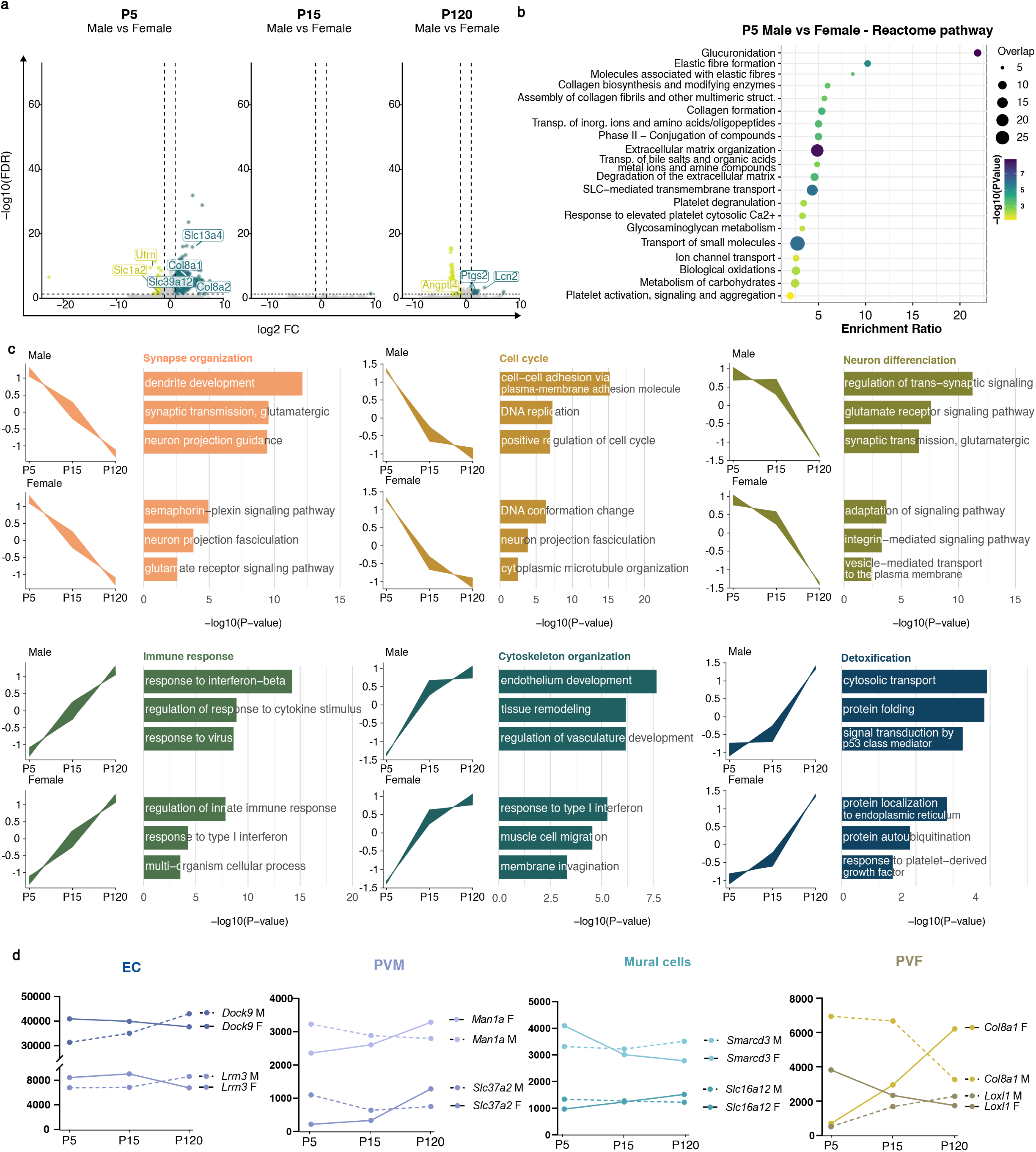
Sex-dependency of the postnatal transcriptome in the GVU. **a.** A volcano plot showing differentially expressed genes (DEGs) in males vs. females on P5 (left), P15 (middle), and P120 (right). Genes that were upregulated in males (relative to females) are shown in green, and genes that were downregulated are shown in yellow. Genes that were neither upregulated nor downregulated are shown in grey. n_P5Male_ = 3, n_P5Female_ = 4, n_P15Male_ = 2, n_P15Female_ = 3, n_P120Male_ = 4, n_P120Female_ = 3. *Ddx3y, Uty, Kdm5d,* and *Eif2s3y* (linked to the Y chromosome) are not illustrated in the volcano plots because they are expressed approximately 1,000-fold higher in male tissues than in female tissues, which would compress the scale and obscure the other differences. **b** A top 20 Reactome pathway analysis comparing males and females on P5. **c.** Expression profile during development, showing genes that were downregulated between P5 and P120 (upper panel) and genes that were upregulated between P5 and P120 (lower panel). Gene sets were clustered by the enriched biological process. **d.** Expression of vascular-cell-specific genes, showing opposite trends in males vs. females from P5 to P120. Dashed lines represent male (M) genes, and solid lines represent female (F) genes.

We next compared the GVU’s transcriptomic developmental trajectories in male vs. female mice from P5 to P120. Six clusters (corresponding to distinct developmental profiles) were found in both males and females (**Fig. 6d; Table S2**). Three downregulated clusters were enriched for biological processes related to *Synapse organization*, *Cell cycle*, and *Neuronal differentiation*. In contrast, three upregulated clusters were associated with *Immune response*, *Cytoskeletal organization*, and *Detoxification* (**Fig. 6c; Table S2)**. Although similar clusters were identified in both males and females, they were not strictly identical: the expression of 107 DEGs changed in opposite developmental directions (**Table S3)**. By focusing on cell-specific or enriched DEGs (expressed in at least 60% of cells of a given type and in fewer than 40% of cells of other types, according to ^63,64^), we identified candidates for the sex-specific maturation of the GVU. For example, *Smarcd3* (BAF60c) promotes the contractile differentiation of VSMCs ^71^ (**Fig. 6d; Table S3**). This gene was progressively downregulated in females and upregulated in males. The same profile was found for *Lox1l1*, which is expressed in PVFs and encodes an enzyme essential for elastin fibre maintenance and the stability and elasticity of arteries ^72^ **(Fig. 6d; Table S3)**. In contrast, *Man1a* and *Slc37a2* (expressed in PVMs) were downregulated progressively in males and upregulated progressively in females (**Fig. 6d; Table S3)**. *Man1a* encodes an enzyme that removes α-linked mannose from high-mannose-type glycans, which are strongly expressed by immature macrophages ^73^. *Slc37a2* has been shown to control macrophage glycolysis and innate activity ^74^.

Lastly, we focused on transcriptomic changes between P5 and P15, which is a crucial developmental window for the GVU (**Fig. 7; Table S1**). Again, DEGs were defined as a log2 FC ≤ -1 or log2FC ≥ 1 and an adjusted p-value ≤ 0.05 (**Fig. 7a; Table S1**). In total, we identified 2237 DEGs between P5 and P15. Of these, 1232 were regulated the same way in the two sexes, while 648 displayed differential expression in females only and 357 displayed differential expression in males only **(Fig. 7b; Table S1)**. To further analyse these changes, we again focused on cell-specific or enriched transcriptomic signatures (i.e. genes expressed in at least 60% of cells of a given type and in fewer than 40% of cells of other types, according to ^63,64^). Although some cell-specific signature genes showed the same trajectory, the sexes differed with regard to the FC; the transcriptional shifts were generally greater in females. This was the case for the PVF-specific genes *Col1a1, Efemp1*, *Lum* and *Cyp1b1* (**Fig. 7c; Table S1**) and the macrophage-specific genes *Cd14* and *C5ar1*. Between P5 and P15, all of these genes were more strongly upregulated in females than in males.

**Figure 7:**
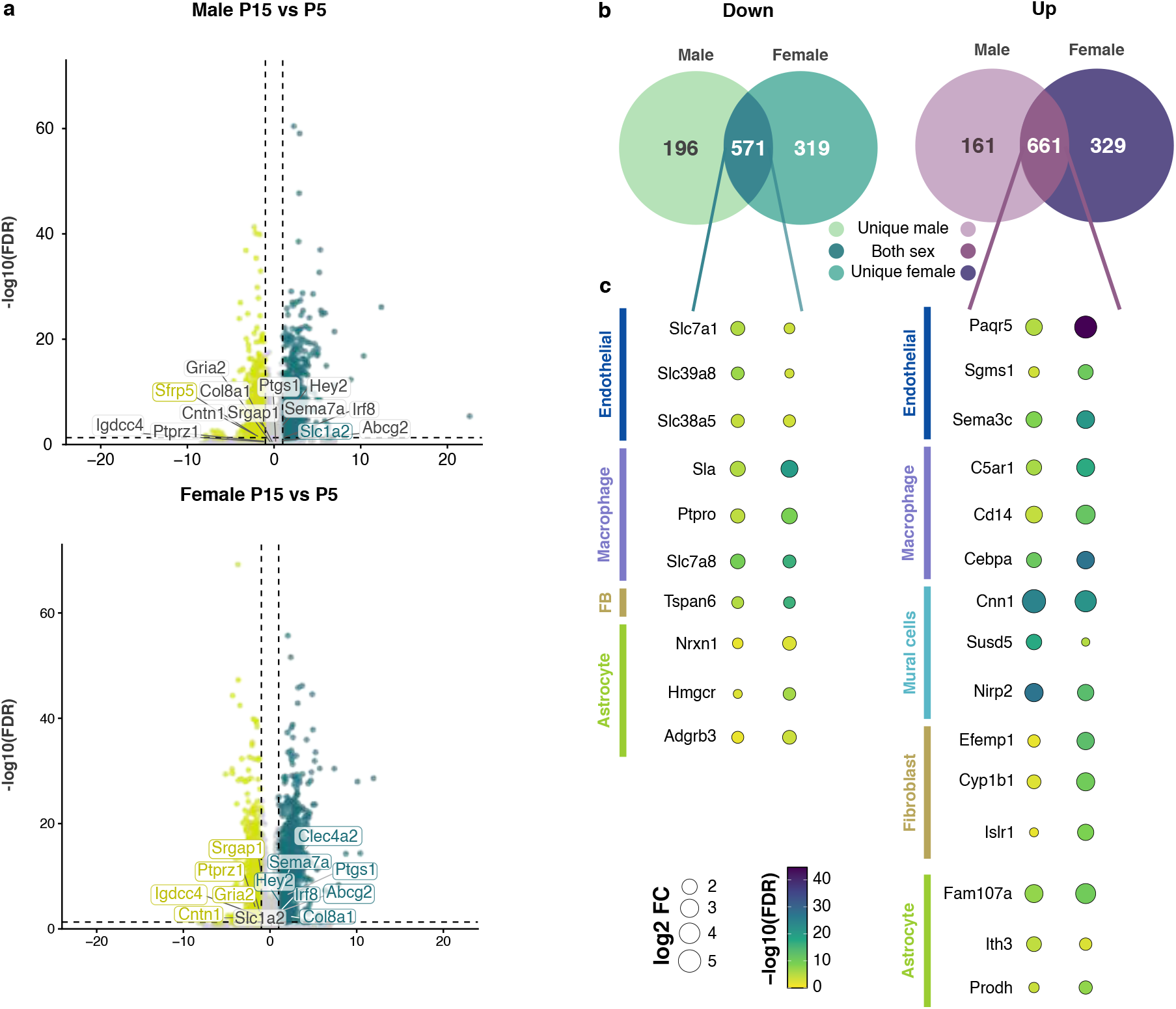
Sex-dependent gene expression trajectories in the GVU. **a** A volcano plot showing DEGs between P15 and P5 in females (left) and in males (right). Genes that were upregulated on P15 (compared with P5) are shown in green, and genes that were downregulated are shown in yellow. Genes that were neither upregulated nor downregulated are shown in grey. **b.** A Venn diagram showing sex-specific and non-sex-specific DEGs on P15 vs. P5. **c.** A dot plot of DEGs in specific vascular cell types, comparing P15 with P5 in males and in females. The dot size is proportional to the fold-change. The statistical tests applied and the exact p-values are given in **Table S1.**

When considering the set of genes regulated differentially between P5 and P15 in females only, all the endothelial genes were found to be upregulated (**Fig. 67; Table S1**). These genes included *Sema7a* (encoding semaphorin 7a, a secreted protein crucial for the development of neuronal circuits in the somatosensory cortex ^75^) (**Fig. 7c; Table S1**). Among the genes expressed mostly by VSMCs, we noted the upregulation from P5 to P15 of *Hey2/CHF1*, a Notch -target gene ^76^ that might mediate arterial cell fate ^77^ (**Fig. 7c; Table S1**). Several of the genes enriched in PVMs were highly upregulated on P15, including *Irf8* (encoding the interferon regulatory factor 8 involved in PVM development and function)^78,79^ (**Fig. 7c; Table S1**). The most prominent change concerned the upregulation of *Clec4a2* (encoding a C-type lectin receptor responsible for the differentiation of monocytes into vascular resident macrophages; FC: 13.08)^80^ (**Fig. 7c**). Lastly, several genes enriched in astrocytes were up- or downregulated (**Fig. 7c; Table S1**). Strikingly, the majority of these were downregulated on P15, and all were related to the development of neuronal circuits: *Ptprz1* ^81^; *Nnat* ^82^; *Srgap1*^83^; *Cntn1* ^84^; *Astn1* ^85^; *Cspg5* ^86^; *Igdcc4* ^87^; *Gria2* ^88^ (**Fig. 7c; Table S1**). The genes regulated between P5 and P15 in males included *Slc1a2*, which encodes the glutamate transporter 1) (**Fig. 7c; Table S1**). Male expression of the astrocyte-specific *Sfrp5* (encoding the secreted frizzled-related sequence protein 5, an inhibitor of Wnt) fell to female levels on P15 (**Fig. 7c; Table S1**).

Our results show that GVUs from male mice and those from female mice have the same overall transcriptomic developmental profiles, with the downregulation of cell cycle processes and synaptic maturation and the upregulation of vascular properties. However, the early postnatal period on P5 is highly dysmorphic. These early transcriptomic differences are likely to drive the sex-specific dynamics of GVU formation and maturation.

## Discussion

The GVU is a critical interface between the brain and the vascular system. It orchestrates essential homeostatic functions, including BBB integrity, metabolic exchanges, waste clearance, neurovascular coupling, and immune surveillance. GVU dysfunction is a common feature of many neurological disorders. Although sex-dependent differences in neurodevelopmental trajectories and disease prevalence are well documented, their impact on GVU assembly and function has not been extensively explored. Here, we compared the GVUs in male and female mice on P5 and P15 (the time window for GVU maturation) and in adulthood. Our data demonstrate that males and females differ with regard to the postnatal GVU’s anatomy, molecular repertoire, and function.

Males showed a transiently higher capillary density in the somatosensory cortex, which was correlated with a higher level of *Vegfa* expression in astrocytes and a higher number of tip cells. These results are consistent with a previous report of higher P14 levels of VEGF-A (measured with an ELISA) in males than in females ^89^. Thus, the astrocyte-driven angiogenic dynamics in the somatosensory cortex are much more active in males on P15. Somatosensory astrocyte density was also transiently elevated in males on P15. In a recent study, this effect was linked to a lower level of astrocyte phagocytosis by microglia in males and specifically on P15 ^23^. Notably, the male vs. female difference in P15 astrocyte and capillary density had no impact on the level of astrocyte perivascular coverage and the distance between astrocyte soma and vessels; this finding suggests that the development of astrocytes and the development of blood vessels are tightly coupled.

While the extent of astrocyte perivascular coverage was equivalent in males and females, the PvAP Aqp4 level was transiently higher in males on P15 and P30; this suggests that the acquisition of perivascular water homeostasis is subject to sexual dimorphism. The same trend was observed previously in the hippocampus on P12 by Arladan and colleagues ^90^. In contrast, Ferreira-Rodrigues and colleagues ^91^ reported no significant difference on P14 in the somatosensory cortex, although there was a trend toward greater Aqp4 coverage in males. These discrepancies are probably due to differences in sampling size, with a large-field view in our study vs. a view of individual capillary segments in ^91^. We identified *Aqp4* mRNA splicing and the production of M1 and M23 Aqp4 isoforms as the mechanism that potentially underlies the sex difference in perivascular Aqp4 levels. A recent characterization of cortical astrocyte polysomal mRNAs demonstrated that astrocytes in males mature more quickly than astrocytes in females (by P14, specifically) ^24^. It would therefore be interesting to look for further sex differences in the development of PvAPs.

We have shown previously that the postnatal contractile maturation of VSMCs in mice and humans is correlated with a progressive increase in CBF ^50^. We have also shown that CBF develops earlier in the lateral cortex than in dorsal cortical regions ^50^. Here, we observed the same dorsolateral difference on P15. Furthermore, dorsal CBF was higher in females than in males. Taken as a whole, these data confirm the differences in the time course of cortical CBF development and demonstrate that P15 females are more mature than P15 males. Another element that might contribute to this effect is the elevated number of Lyve-1+ PVMs. Indeed the presence of these cells is correlated with a higher CBF ^92^. Together, these differences might be directly involved in neuronal development and function because neurotransmission is highly dependent on an adequate blood supply. The second wave of postnatal brain angiogenesis is tightly coupled to neuronal maturation ^15^ and activity-dependent neurovascular interactions have been demonstrated previously ^93^; hence, sex-dependent neuronal maturation and circuit activity might also contribute to differences in vascular maturation. This could be addressed further by evaluating the development of male and female GVUs in models with pure synaptogenic impairments, e.g. synaptic hearing loss models with a low vascular density in the auditory cortex ^93^. Importantly, our observation of enhanced expression of SMA in human tissues indicates that arterial development and the CBF might also be faster in female infants than in male infants, as already reported by others ^94^. These differences might have a protective effect in females and might account for the greater prevalence of certain neurodevelopmental disorders (e.g. autism spectrum disorder) in males.

We were intrigued to find that from P15 onward, phenotypic sexual dimorphisms of the GVU become apparent in the absence of detectable transcriptomic differences. This observation suggests that sexual maturation of the GVU by P15 is not governed primarily by transcriptional mechanisms.

An additional layer of analysis (e.g. a proteomic comparison of male and female MVs) might reveal further sex-specific differences.

In contrast to P15, the P5 GVU transcriptome showed major male vs. female differences on P5. Our transcriptomic analysis showed than P5 female PvAPs already exhibited mature expression levels of *Slc1a2* (encoding the glutamate transporter GLT-1, one of the most abundant astrocytic proteins in PvAPs) ^95,96^. If this additional amount of mRNA is translated into protein, glutamate clearance by vessel-associated astrocytes might be enhanced in females in the early postnatal stages. Another striking observation in females was the greater downregulation between P5 and P15 of several astrocyte-enriched genes involved in synapse formation, including *Cntn1, Ptprz1, Nnat, Astn1*, and *Cspg5*. This finding might correspond to sex-specific dynamics in the assembly of perivascular synaptic structures ^97^. Notably, perivascular water and glutamate homeostasis and the assembly of the synapses abutting PvAPs are essential for neurovascular coupling, i.e. the coordination between synaptic and vascular activities ^13^. Our data indicate that males and females differ with regard to the development of this essential GVU function.

ECM-related genes (including *Col8a1, Col8a2, Col1a1, Efemp1,* and *Lum*) were upregulated more strongly in female mice than in male mice. These transcriptional differences might influence the maturation and composition of perivascular basement membranes (particularly around large vessels), with potential consequences for arteriolar membrane properties and CBF regulation ^38^. Given that PVFs contribute to PVM recruitment, enhanced PVF maturation in females might also influence dimorphism of the immune compartment ^98^. Consistently with this hypothesis, we observed more Lyve1^+^ PVMs in females on P15, and transcriptomic analyses revealed the sex-specific regulation of macrophage-associated genes: *Cd14* and *C5aR1* were upregulated between P5 and P15 exclusively in females and *Cd163* (a marker of brain-resident PVMs) showed a greater increase in females. PVMs have key roles in CSF drainage and immune surveillance. Lyve1^+^ PVMs have been shown to influence arterial pulsation and vasomotion ^92^. PVMs have been implicated in the pathophysiology of amyloid clearance, hypertension, stroke, and aneurysm ^45,46,92^. Differential PVM recruitment in females might contribute to faster arteriolar maturation, higher CBF, and sex-biased vulnerability to cerebrovascular disorders.

Sex steroid signalling provides a plausible mechanistic framework for these observations. Indeed, cerebral blood vessels express oestrogen receptors (ERα, ERβ, and GPER1), androgen receptors, and enzymes (such as aromatase) that convert testosterone to oestrogens locally ^61^. Circulating sex steroids can access the brain through the BBB ^60^, and animal studies have evidenced the sex steroids’ ability to modulate not only endothelial permeability ^10^ and vascular tone ^99–101^ but also astrocyte growth ^102^, calcium signalling ^103–105^, and Aqp4 expression ^106^. Macrophages and fibroblasts are also responsive to sex steroids, and oestrogens and progesterone regulate collagen expression - providing a potential link between hormone signalling and ECM remodelling ^107^. A recent report showed that oestrogens also control microglial phagocytic capacities on P15 in mice and thus influence somatosensory astrocyte density and synapse remodelling ^23^. Our present results show that oestradiol delivered to females at birth (mimicking the male perinatal surge of testosterone converted into neural oestradiol) masculinizes the female vascular network by diminishing the SMA+ vessel density, stimulating angiogenesis, and increasing capillary density. In contrast to Duffy et al., we did not detect an effect on astrocyte density ^23^. This discrepancy might be related to a dose-dependent effect, as the mice in the present study received only 20 µg oestradiol benzoate on P1 - a dose shown previously to induce masculinizing effects in sexually differentiated brain regions in mice ^108,109^.

In summary, we comprehensively characterized sex-specific developmental trajectories within the GVU. Most of the cellular compartments of this vascular interface exhibited sexually dimorphic maturation programs that were controlled (at least in part) by oestrogen signalling. Our findings underscore the importance of incorporating sex as a biological variable in cerebrovascular research and establish a framework for investigating how early-life GVU dimorphism might shape long-term susceptibility to neurological disease.

### Principal limitations of the study

The present study had a number of limitations. Firstly, we used bulk RNA sequencing of purified MVs to compare male and female gliovascular transcriptomes. Consequently, for genes that are not specific markers of a given cell type, it was not possible to determine whether the observed transcriptional differences reflected changes within a particular cell population or differences in cellular composition. Single-cell transcriptomic approaches provide greater resolution and would enable a more precise characterization of sex-dependent transcriptional differences. Secondly, given the dynamic nature of postnatal brain development, it is still not clear whether the observed sex differences reflect developmental delays, transient shifts, or genuinely distinct developmental trajectories. Future longitudinal studies (including analyses across closer developmental stages) might be needed to distinguish between these possibilities. Lastly, vascular geometric variables may be influenced by fixation and tissue sectioning. *In vivo* vascular imaging could provide a more comprehensive characterization of gliovascular sexual dimorphism during postnatal brain development.

## Methods

### Reagents

All reagents are described in the Supplementary Data **(Table S4)**.

### Mice

C57Bl/6j mice were provided by Janvier Labs. Tg(Aldh1l1-eGFP) OFC789Gsat/Mmucd (Aldh1l1:eGFP) mice were obtained from Nathaniel Heintz (Rockefeller University, NY, USA) and kept under pathogen-free conditions. All animal experiments were conducted in accordance with European Parliament Directive 2010/63/EU and were approved by an institutional animal care and use committee and the French Ministry of Higher Education and Research (reference numbers: 37351, 48496, 35402 and 51249). The mice used for immunostaining and P5 and P15 transcriptome analysis are included as controls in another project (manuscript in preparation) and received tap water containing 0.001% DMSO v/v. This concentration has no known baseline effects on brain development and function^110,111^. For all experiments, males and females of the same litter were compared. Exceptions to this rule are indicated in **Table S5**.

### Oestradiol treatment

Animals received a subcutaneous injection of 20 µg β-oestradiol benzoate (Sigma-Aldrich, E8515) in 20 µL of sesame oil on P1. This dose was shown previously to be sufficient to induce masculinizing effects in sexually differentiated brain regions in mice^108,109^. Mice were left in their parent cage until P15, when they were euthanized and the brains were harvested for analysis.

### Immunohistofluorescence: imaging and analysis

Mice were perfused transcardially with 4% paraformaldehyde (PFA), and brains were post-fixed in 4% PFA overnight at 4°C. After dehydration in 30% sucrose, brains were cut into 60-μm-thick sections using a Leitz (1400) cryomicrotome and kept at -20°C in storage solution (PBS/30% glycerol/30% ethylene glycol). Brain sections were rinsed in PBS for 15 min three times and incubated in a blocking solution (PBS/0.2% gelatine/0.5% Triton X-100) for 1 h at room temperature (RT). Sections were incubated overnight at 4°C with primary antibodies diluted in the blocking solution. After three 15-min rinses in PBS, the sections were incubated for 2 h at RT with secondary antibodies (Supplementary Table 4) and Hoechst reagent (Supplementary Table 4), rinsed in PBS, and mounted in Fluoromount-G^TM^ (Southern Biotech, Birmingham, AL, USA).

Images were acquired using a wide-field Axio Zoom V16 microscope and its Apotome V2 module (Zeiss) or a spinning disk widefield confocal scanner unit (Zeiss and Nikon), both of which were equipped with an sCMOS Hamamatsu 2304x2304 camera or a confocal LSM 980 (Zeiss).

The CD31 network was analysed through the three-dimensional (3D) reconstruction of cortical blood vessels using Imaris software (Oxford Instruments, Oxford, UK, version 10.1). The vessel length, number of segments, segment length, branch angles, and segment straightness of CD31 blood vessels were quantified using the “Surface” and “Filament” tools. The vessel length, number of segments, and segment length were normalised against the somatosensory cortical volume. Tip cells were analysed using QuPath software (version 0.6.0)^112^. Raw values were normalised against the somatosensory cortical volume. Pericyte coverage was analysed using Fiji/ImageJ software (version 1.54i)^113^. Image segmentation was used creating two masks: one for IB4-stained or CD31-immunolabeled blood vessels (using Yen filter) and one for CD13-immunolabeled pericytes (using the Otsu filter). The filters were adjusted manually to match the labelling and optimize the signal-to-noise ratio. For each marker, a region of interest was created. The “AND” function was used to identify areas of colocalisation. Raw area values were normalised against the total local vascular area. Aqp4 and Aqp4ex were analysed in the same way. Two masks were created, one for IB4-stained or CD31-immunolabeled blood vessels (using the Yen filter) and one for Aqp4/Aqp4ex (also using the Yen filter). Raw area values were normalised against the total local vascular area. Aldh1l1-eGFP coverage was analysed using Imaris software (version 10.1, Bitplane, Zürich, Suisse). 3D reconstructions of cortical blood vessels and astrocyte-GFP were created using the Surface tool. The contacts between GFP-expressing astrocytes and blood vessels were reconstructed using the “Surface-Surface contact area” Imaris Xtension, which generates a one-voxel-thick, unsmoothed surface on the vessel. All 3D-rendered surfaces were used by the software to calculate the surface area of the elements. We analysed Sox9-positive cells using a custom-developed macro for Fiji/ImageJ (version 1.54i). Sox9-positive cells were detected using a 2D-stitched version of the *cellpose* algorithm (https://github.com/orion-cirb/Orion_Macros.git) with the exception of masculinization experiments, in which cells were counted manually (to avoid mistakenly counting erythroid cells, as the mice were not perfused). A custom model was trained through the workflow graphical user interface available in *cellpose* ^25^. Only cells with a volume of at least 50 µm³ were detected. Raw values were normalised against the somatosensory cortical volume. Distances between Sox9^+^ astrocyte somas and blood vessels were analysed using Imaris software. Two surfaces were created, one for IB4-stained or CD31-immunolabeled blood vessels and one for Sox9^+^ astrocyte somas using the “Surface” tool. Next, the filter “Shortest distance to surface” was applied with a threshold of 0.8 µm. Sox9^+^ astrocyte somas within 0.8 µm of a blood vessel were considered to be perivascular astrocyte somas. Raw values were normalised against the somatosensory cortical volume. The proportion of perivascular astrocyte somas was obtained by dividing the number of perivascular astrocyte somas by the total number of Sox9+ astrocyte somas.

The total number of penetrating blood vessels and the number of penetrating blood vessels with at least one PVM were counted with Fiji. The ratio between the number of blood vessels with one PVM and the total number of penetrating blood vessels was calculated. For calculation of CD206 and Lyve-1 density, images were analysed with QuPath software. The “Cell detection” plug-in was used, and the parameters were set as follows for CD206 and Lyve-1, respectively: requested pixel size: 2 µm and 3 µm; background radius: 10 µm and 20 µm; median: 1 µm for both; sigma: 3 µm for both, min: 100 µm² for both, max: 1000 µm² for both, threshold: 200 µm and 100 µm. To determine PVM categories, we referred to the eGFP/Cy3 range value; this is below 100 for CD206^+^ Lyve-1^-^ PVMs, equal to or greater than 100 for CD206^+^ Lyve-1^+^ PVMs, and below 550 for CD206^-^ Lyve-1^+^ PVMs. Raw values were normalised against the volume of the field of view.

### Fluorescence *in situ* hybridization and analysis

FISH was performed on floating fixed brain sections, using the v2 Multiplex RNAscope technique (Advanced Cell Diagnostics, Inc., Newark, CA, USA). After the FISH, blood vessels were immunostained for CD31, as described previously ^50^. Images were acquired using a wide-field Axio Zoom V16 microscope and its Apotome V2 module (Zeiss), equipped with an sCMOS Hamamatsu 2304x2304 camera. The *Col1a1* signal was analysed using QuPath software (version 0.6.0). The “Cell detection” plug-in was used, and the parameters were set as follows for the *Col1a1* signal: requested pixel size: 2 µm; background radius: 10 µm; median: 1 µm; sigma: 3 µm, min: 100 µm², max: 3000 µm², threshold: 100 µm. The *Vegfa* signal was analysed using Fiji/ImageJ (version 1.54i). Image segmentation was used creating a mask using Yen filter. The filters were automatically adjusted to match the labelling and optimize the signal-to-noise ratio. Raw values were normalised against the volume of the field of view.

### Immunohistochemical analysis of human tissue samples

The cortical sections studied here are part of the *Hôpitaux Universitaires de l’Est Parisien –Neuropathologie du développement* brain collection (Paris, France; biobank identification number: BB-0033-00082). Informed consent was obtained for the brain autopsies and histological examinations. Foetal brains were obtained from spontaneous or medical abortions. The foetuses did not display any significant brain disorders or diseases. Postnatal brain samples were obtained from children who died from non-pathological causes. The following samples were analysed: female, 17 weeks of amenorrhea (WA); female, 23 WA; male, 30 WA; male, 32 WA; female, 40 WA; female, 3 weeks old; male, 1 month; male, 2 months; female, 8 months; male, 1 year; male, 3.5 years; male, 4 years; male, 10 years; male, 11 years; female, 12 years; female, 13 years (n=2); male. One slice per sample was analysed. After removal, the brains were fixed with formalin for 5 to 12 weeks. A macroscopic analysis enabled the samples to be selected and processed for histological analysis (paraffin embedding, preparation of 7-µm-thick slices, and staining with hematein reagent). Coronal slices (including the temporal telencephalic parenchyma and the hippocampus) were deparaffinized and unmasked in citrate buffer (pH 6.0). Expression of SMA was detected using the Bond Polymer Refine Detection kit (Leica, Newcastle, UK) and processed on an automated immunostaining system (the Leica BOND RX for MYH11, and the Leica BOND III for SMA). Images were acquired using a slide scanner (Lamina, Perkin Elmer).

The images of the stained samples were analysed using QuPath software ^112^. For each sample, a QuPath “pixel classifier” was trained to discriminate between 3, 3’-diaminobenzidine-positive spots and the background. This “classifier” consisted of an artificial neural network based on four features: a Gaussian filter to select the intensity, and three structure tensor eigenvalues to favour thin, elongated objects. To train the classifier, we manually annotated spots and background area on one image per developmental stage. When the results were visually satisfactory, the trained pixel classifier was used to detect positive spots in manually defined regions of interest.

### Brain clearing: imaging, and analysis

Brains were post-fixed in 4% PFA for 24 h at 4°C and cleared as described in ^114^. The samples were first dehydrated with aqueous methanol series (20%, 40%, 60%, 80%, and then 100% twice, for 1 h each) at RT and then incubated in 66% dichloromethane (Sigma-Aldrich)/ 33% methanol overnight. After two washes in 100% methanol, brains were incubated in 5% H_2_O_2_/methanol overnight at RT, and rehydrated with a methanol/H_2_O series (80%, 60%, 40%, and 20%; 1 h each). Before immunostaining, brains were permeabilized for 2 × 1 h at RT in 0.2% Triton X-100/PBS, for 24 hr at 37°C in 0.16% Triton X-100/2.3% glycine/20% DMSO/PBS, and lastly for 2 days at 37°C in 0.16% Triton X-100/6% donkey serum/10% DMSO/PBS. Brains were incubated for 3 days at 37°C with primary antibody diluted in a 0.2 Tween-20/1% heparin/3% donkey serum/5% DMSO/PBS solution, washed five times for 24 h at 37°C in 0.2% Tween-20/1% heparin/PBS solution, incubated for 3 days at 37°C with secondary antibody diluted in a 0.2% Tween-20/1% heparin/3% donkey serum/PBS solution, and washed another five times. The samples were then dehydrated again with a methanol/H_2_O series (20%, 40%, 60%, 80%, and 100% for 1 h each, and then 100% overnight) at RT. On the following day, brains were incubated for 3 h in 66% dichloromethane/33% methanol and then twice for 15 min at RT in 100% dichloromethane and, lastly, cleared overnight in dibenzyl ether. Cleared brains were imaged using a light sheet microscope and Inspector pro software (Lavision Biotec GmbH, Bielefeld, Germany). 3D reconstructions of the somatosensory cortex were produced using Imaris software. The length and number of branch points on SMA-immunolabeled brain vessels were quantified using the Imaris “Surface” and “Filament” tools.

### MRI of CBF

MRI was performed on a 7T Pharmascan system (Bruker®, Germany) equipped with volume transmit and surface receive coils. To prevent motion during the acquisition, animals were anesthetized using one of two protocols: isoflurane or medetomidine. For the isoflurane group, anesthesia was induced with 5% isoflurane delivered in a 30% O2/70% N2O gas mixture, and mice were maintained under 2-3% isoflurane throughout the acquisition, with respiratory rate regulated at 70-100 breath per minute by adjusting the isoflurane concentration. For the medetomidine group, anesthesia was induced with 5% isoflurane delivered in a 30% O□/70% N□O gas mixture until loss of the righting reflex and a marked decrease in respiratory rate were observed. Isoflurane was then reduced to 3% in the same gas mixture, and mice received a single subcutaneous bolus injection of medetomidine (Domitor, Vetoquinol, Lure, France) at 55 µg/kg (40 µl). Isoflurane was immediately lowered to 1.5%, and a continuous subcutaneous infusion of medetomidine, prepared at the same dilution as the bolus solution, was started at 135 µg/kg/h using a syringe pump (AL-1000, World Precision Instruments). Isoflurane concentration was then progressively decreased by 0.25% every 2 min until complete discontinuation of isoflurane, O□, and N□O. Mice were maintained under spontaneous ventilation throughout the sedation period. Sedation was maintained for the duration of the imaging session by continuous subcutaneous infusion of medetomidine alone, at the rate indicated above. For both protocols, core body temperature maintained at 37 °C using a heating system. The total acquisition time did not exceed 20 min. Before the ASL sequence, a fast T2 anatomic sequence was acquired to allow precise targeting of the slice of interest. Common anatomic features allowed the targeting of the intended slice (bregma +1 mm) in all groups. T2-weighted images were acquired using an 18-slice multi-echo sequence (TE/TR, 39.99 ms/3500 ms; 4 averages). The ASL sequence was performed to measure brain perfusion at bregma +1 mm (TE/TR, 79/10224.65 ms, 1 average). The built-in ASL sequence on the Bruker MRI scanner was used at a 0.156 x 0.156 mm planar resolution in a single, 1-mm-thick slice (matrix size: 128 x 128). The built-in ASL-calculating macros from Bruker were used for CBF estimations. A one-voxel (0.156 x 0.156 mm) Gaussian filter was applied to the resulting map, in order to smooth the voxel-wise calculations. CBF values were measured in the right and left cortices and averaged for each subject before analysis. Measurements on CBF maps were extracted using FiJi ^113^.

### RT-qPCR

RNA was extracted from whole brain using the Rneasy Lipid tissue mini kit (Qiagen, Hilden, Germany). cDNA was then generated using the Superscript™ III Reverse Transcriptase kit (Thermofisher, Waltham, Massachusetts, USA). cDNA levels were measured using the droplet digital PCR (ddPCR) system (Bio-Rad Hercules, CA, USA). Briefly, cDNA, EvaGreen supermix (Bio-Rad) and primers were distributed into approximately 10,000 to 20,000 droplets. Nucleic acids were then PCR-amplified in a thermal cycler (according to the manufacturer’s instructions) and read as the number of positive and negative droplets with a QX200 Droplet Digital PCR System (Bio-Rad). The ratio for each tested gene was normalised against the total number of positive droplets against *Gapdh*.

### MV purification

MVs with associated PvAPs were purified from cortices without meninges, as described previously ^115^. Briefly, cortices were dissected, rolled on absorbent paper (in order to remove the meninges), and resuspended in Hanks’ balanced salt solution (HBSS)/HEPES, using an automated Dounce homogenizer. After an initial centrifugation at 2000 g for 10□min, the pellet was resuspended in HBSS/18% Dextran and centrifuged at 3000 g for 15□min, to separate the myelin from the vessels. This new pellet contained the cortical vessels with attached PvAPs and was resuspended in HBSS/1% bovine serum albumin. The eluate was then filtered on a 20□μm mesh filter. The retained MVs with attached PvAPs consisted of arterioles, venules, and capillaries. MVs were stored at -80°C before mRNA extraction.

### RNA sequencing and analysis

This work benefited from access to the equipment and services at the iGenSeq genotyping and sequencing core facility at the Institut du Cerveau (Paris, France). Total mRNA was extracted from purified cortical MVs using the RNeasy Lipid Tissue Kit (Qiagen). We prepared mRNA libraries according to the manufacturer’s instructions (Stranded mRNA Prep kit, Illumina, San Diego, CA, USA) and sequenced on an Illumina Novaseq 6000 with an S1 (200 cycle) cartridge (2x1600 million 100 base reads, corresponding to 2x50 million reads per sample after demultiplexing). RNA-Seq data were analysed with Genoplice (www.genosplice.com). Sequencing data quality, read distribution (e.g., for potential ribosomal contamination), inner distance size estimation, gene body coverage, and strand specificity were determined using FastQC v0.11.2, Picard-Tools v1.119, Samtools v1.0^116^, and RSeQC v2.3.9^117^. Reads were mapped using STAR v2.7.5a ^118^ on the mouse mm39 genome assembly, and the read count was determined using featureCount from SubRead v1.5.0 ^119^ and the Human FAST DB v2022_1 annotations. Gene expression was estimated as described previously ^120^. Only genes expressed in at least one of the two compared conditions and covered by enough uniquely mapped reads were analysed further. Genes were considered to be expressed if their fragments per kb per million mapped fragments value was greater than that of 98% of the intergenic regions (i.e. the background). The proportion of uniquely mapped reads had to be at least 50%. We analysed gene-level expression using DESeq2^121^. Genes were considered to be differentially expressed when the FC was ≥ 1.5 and the adjusted p-value was ≤ 0.05. Pathway enrichment analyses and gene set enrichment analyses^122^ were performed using WebGestalt (version 0.4.4)^119^ for all regulated genes and by merging results for upregulated genes only and downregulated genes only. The threshold for statistical significance was set to p≤ 0.05. Deconvolution analysis was performed using GEDIT ^123^. Cell population signatures were defined using single-cell RNA-Seq data from GEO datasets GSE99058 and GSE98816 ^124,125^. Gene clustering analyses were performed on raw count data, using clust (version 1.18.0 ^126^) with the following parameters: -t 3 -q3s 3 -n 1000. Over-representation analyses (based on biological process terms from the Gene Ontology knowledge base) were performed using WebGestaltR (version 0.4.6)^127^ and R (version 4.3.2) (https://www.R-project.org). The RNA sequence gene expression data and raw fastq files are available on the GEO repository (www.ncbi.nlm.nih.gov/geo/) under the accession number GSE325705.

### Aqp4 M1 and M23 isoforms quantification

Proportions of AQP4 M1 and M23 transcripts were estimated from a published cortical astrocyte polysomal dataset GSE192490 ^24^ based on the number of reads supporting the exon 5–exon 7 junction that is specific to M1 transcripts (chr18:15533059–15536657) and the exon 6–exon 7 junction that is specific to M23 transcripts (chr18:15533059–15535716). Differences in transcript proportions between experimental conditions (sex or developmental stage) were assessed using Welch’s t-test. The threshold for statistical significance was set to p≤ 0.05.

### Statistical tests

All statistical analyses are summarized in **Source Data Table S5**. Data were expressed as the mean ± standard deviation (SD). All statistical analyses were performed using GraphPad Prism software (GraphPad Software LLC, Boston, MA, USA). For each dataset, normality was tested using the Shapiro-Wilk test. Statistically significant differences between groups were assessed in a Mann-Whitney test, an unpaired, two-tailed Student’s t-test, or (for CBF measurements) a one-tailed t-test. The chi-squared test was used to compare percentages of astrocyte or pericyte coverage around blood vessels. The threshold for statistical significance was set to p-adjusted < 0.05.

## Acknowledgements

We are grateful to the donors and benefactors who support the charities and charitable foundations cited below, and we thank Rachel Ajzen and Léon Iagolnitzer for their generous help. We also thank the administrative services at the CIRB and the Collège de France for their continued support. We thank Audrey Chagnot, Axel Montagne, and Julie Siegenthaler for helpful discussions.

## Author contributions

LL, MAM, TH, RAP, ML and BDP performed experiments; LL, MAM, TH, ML, ACB and BDP analysed data; MR, LN, SMK, DV, LL, BDP, ACB and MCS conceived experiments; BDP and LL assembled figures; MBL provided samples; BDP and MCS wrote the manuscript.

---

**Supplementary Figure 1:**
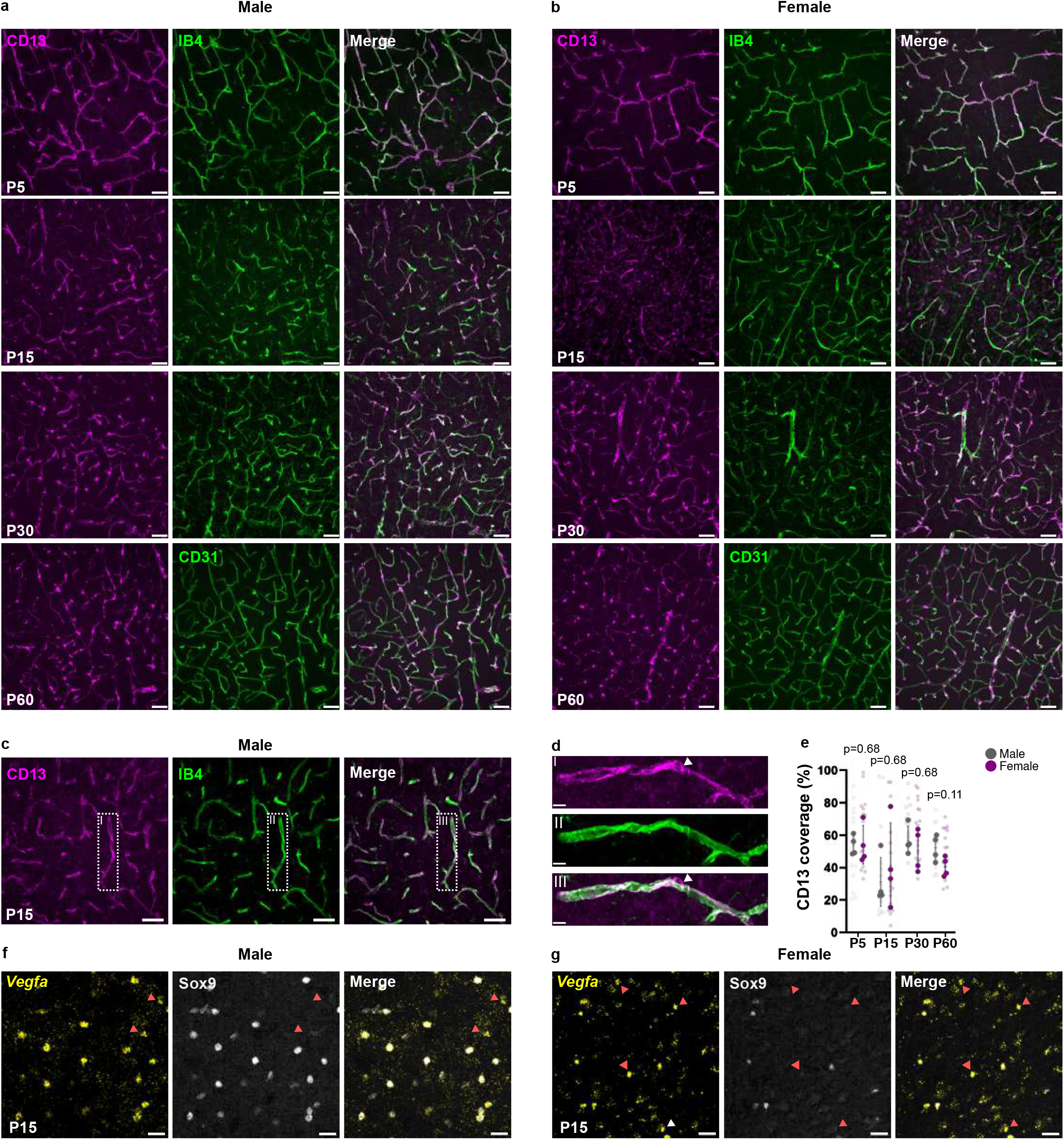
All images for. Figure 1h-j **a-b.** Representative spinning-disk images of pericytes immunolabeled for CD13 (in magenta) in somatosensory cortex sections on P5, P15, P30 and P60 in males and in females. Blood vessels are stained with isolectin-B4 (IB4) (in green) from P5 to P30 and immunolabeled for CD31 (in green) on P60. Scale bar: 50 µm. **c.** Representative spinning-disk images of pericytes immunolabeled for CD13 (in magenta) in somatosensory cortex sections on P15. Blood vessels are stained with IB4 (in green) in males and females. Scale bars: 50 µm. **d.** An enlarged view of a pericyte. The white arrow head indicates a pericyte soma. Scale bar: 10 µm. **e.** Quantification of CD13 perivascular coverage. n= 4 for each sex and each stage. Dark dots represent the mean value for each mouse. Light dots represent values for each analysed image (4 per mouse). n_P5 male_ = 4, n_P5 female_ = 3, n_P15 male_ = 4, n_P15 female_ = 4, n_P30 male_ = 4, n_P30 female_ = 4, n_P60 male_ = 4, n_P60 female_ = 4. The data are presented as the mean ± SD and were analysed using a chi^2^ test. **f.** Representative spinning-disk images of *Vegfa* mRNA detected by FISH (in yellow) in somatosensory cortex sections on P15. Astrocyte somas are immunolabelled for Sox9. In female, red arrow heads indicate Vegfa^+^ Sox9^-^ somas. Scale bars: 50 µm. The raw data are given in **Source Data Table S5**.

**Supplementary Figure 2:**
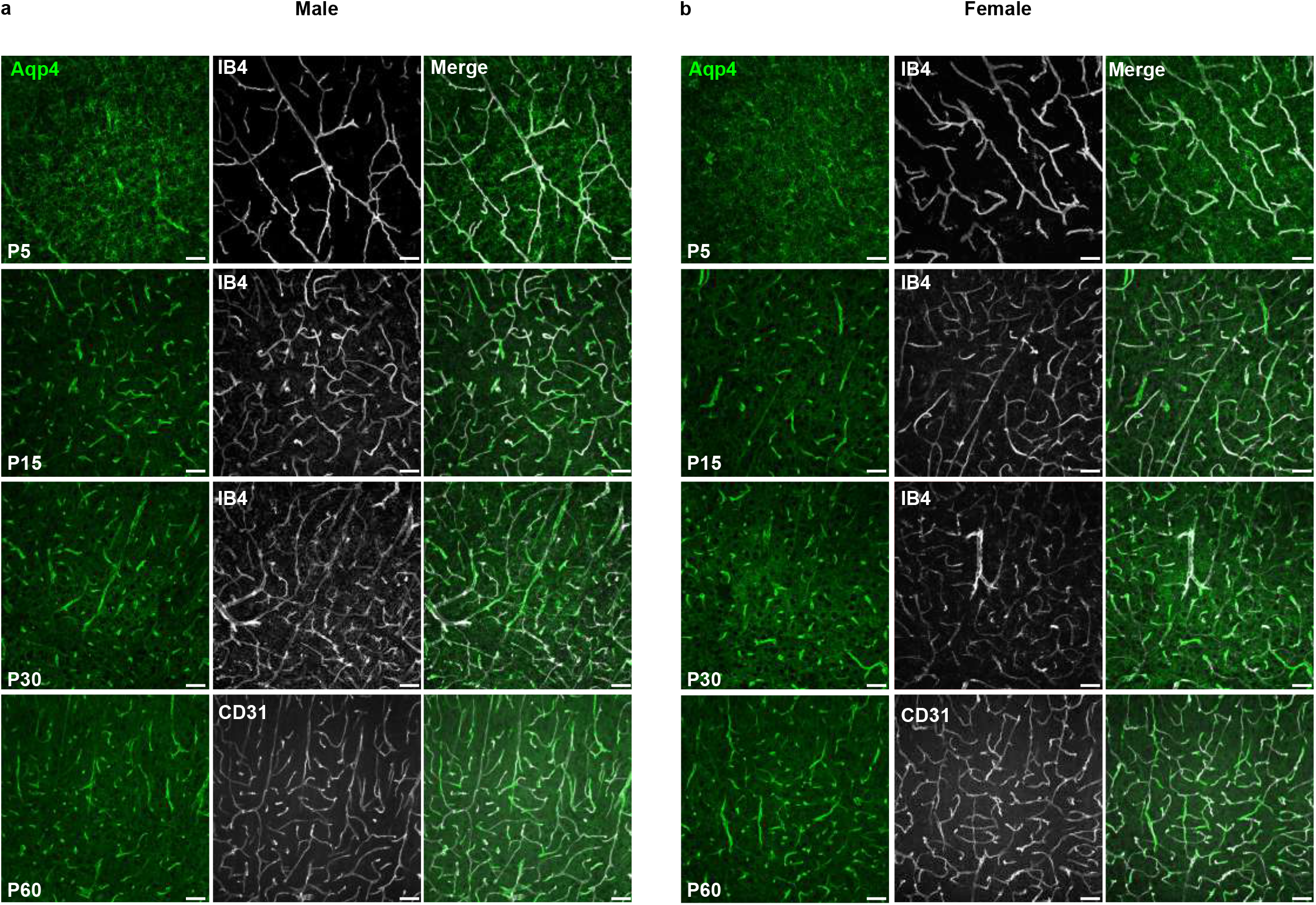
All the images for. Figure 2i **a-b.** Representative spinning-disk images of astrocyte perivascular processes immunolabeled for Aqp4 (in green) in somatosensory cortex sections on P5, P15, P30 and P60 in males and in females. Blood vessels are stained with isolectin—B4 (IB4, in grey) from P5 to P30 and with CD31 (in grey) on P60. Scale bar: 50 µm.

**Supplementary Figure 3:**
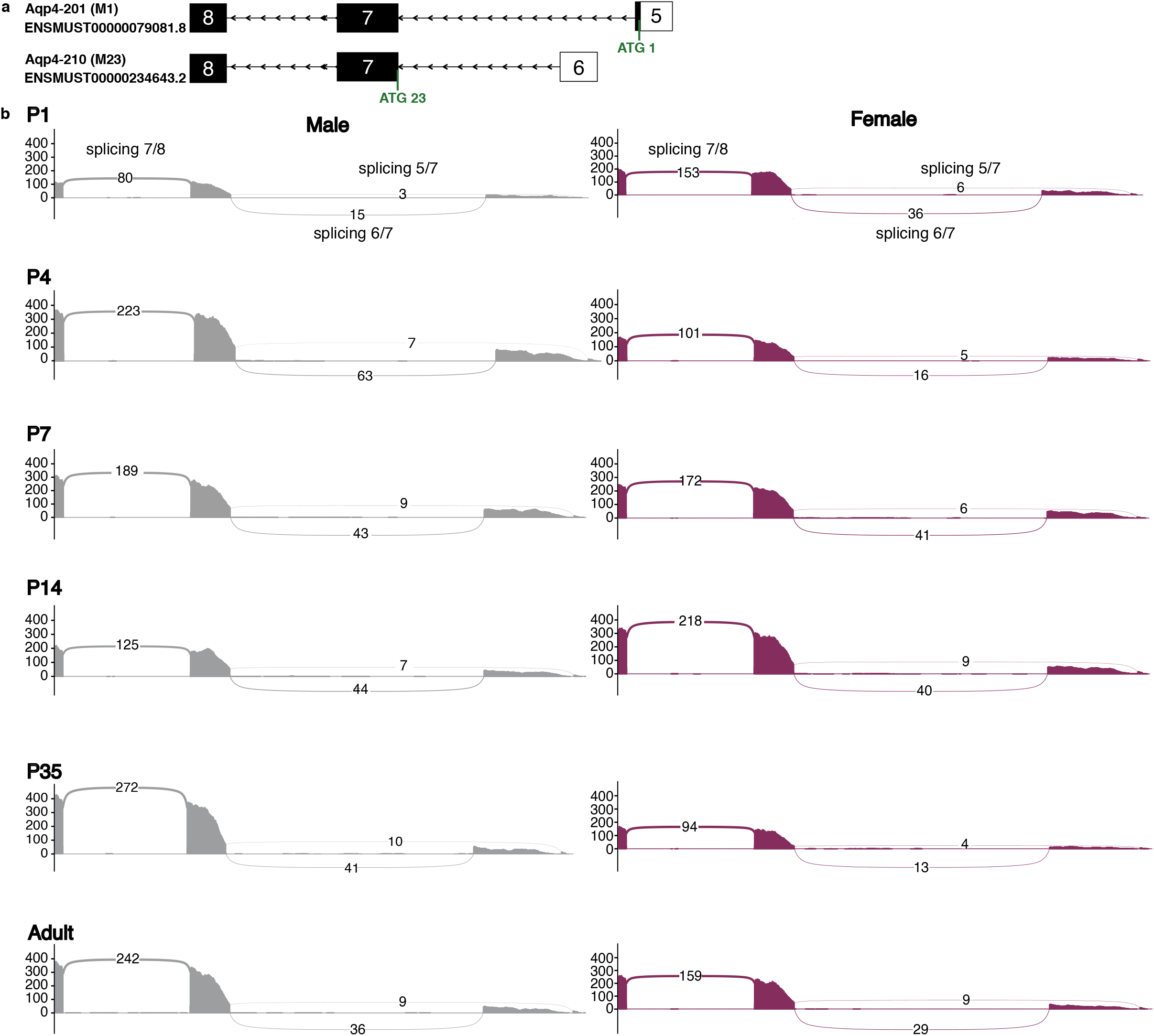
Bioinformatic analysis of Aqp4 M1 and M23 expression in cortical astrocytes. **a.** Exon-intron organisation of *Aqp4M1 and M23.* Exons are indicated by boxes, in black for the coding sequences and in white for uncoding sequences. ATG codons encoding alternative methionine are indicated. **b.** Graphical representations of the M23 and M1 splicing event were generated as sashimi plots using ggsashimi ^128^ over the hg38 genomic region chr18:15531228–15536787. Junction read counts are displayed using the mean aggregation over the samples. The raw data are given in **Source Data Table S5**.

**Supplementary Figure 4:**
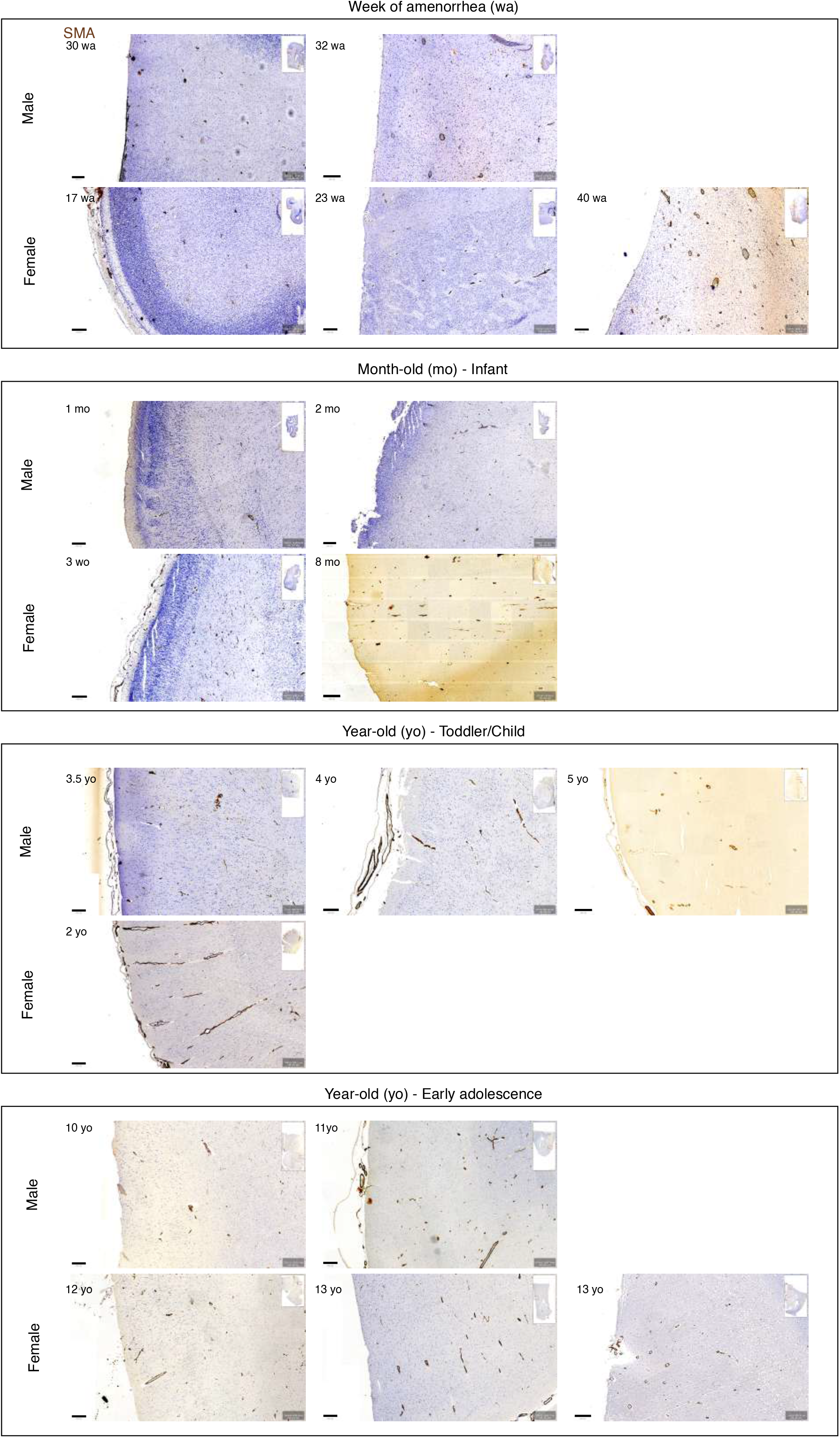
All the images for. Figure 4j **(SMA immunolabelling on human cortex)** Immunohistochemical analysis of smooth muscle actin (SMA) expression in the developing cortex in human males and females (WA: week of amenorrhea, mo: months old, yo: years old). Scale bar: 250 µm.

**Supplementary Figure 5:**
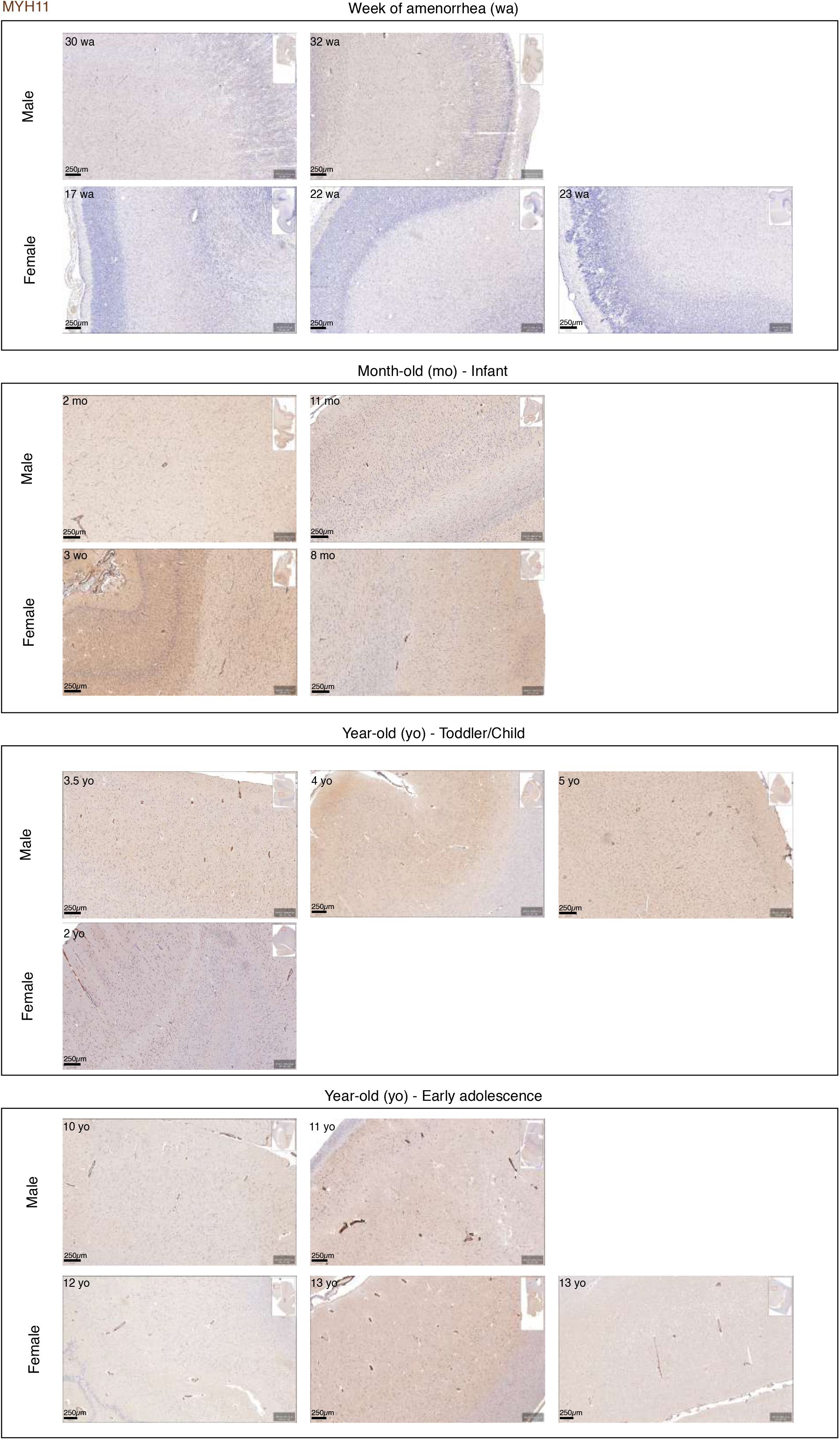
All the images for. Figure 4j **(MYH11 immunolabelling on human cortex)** Immunohistochemical analysis of Myosin 11 (MYH11) expression in the developing cortex in human males and females (WA: week of amenorrhea, mo: months old, yo: years old). Scale bar: 250 µm.

**Supplementary Figure 6:**
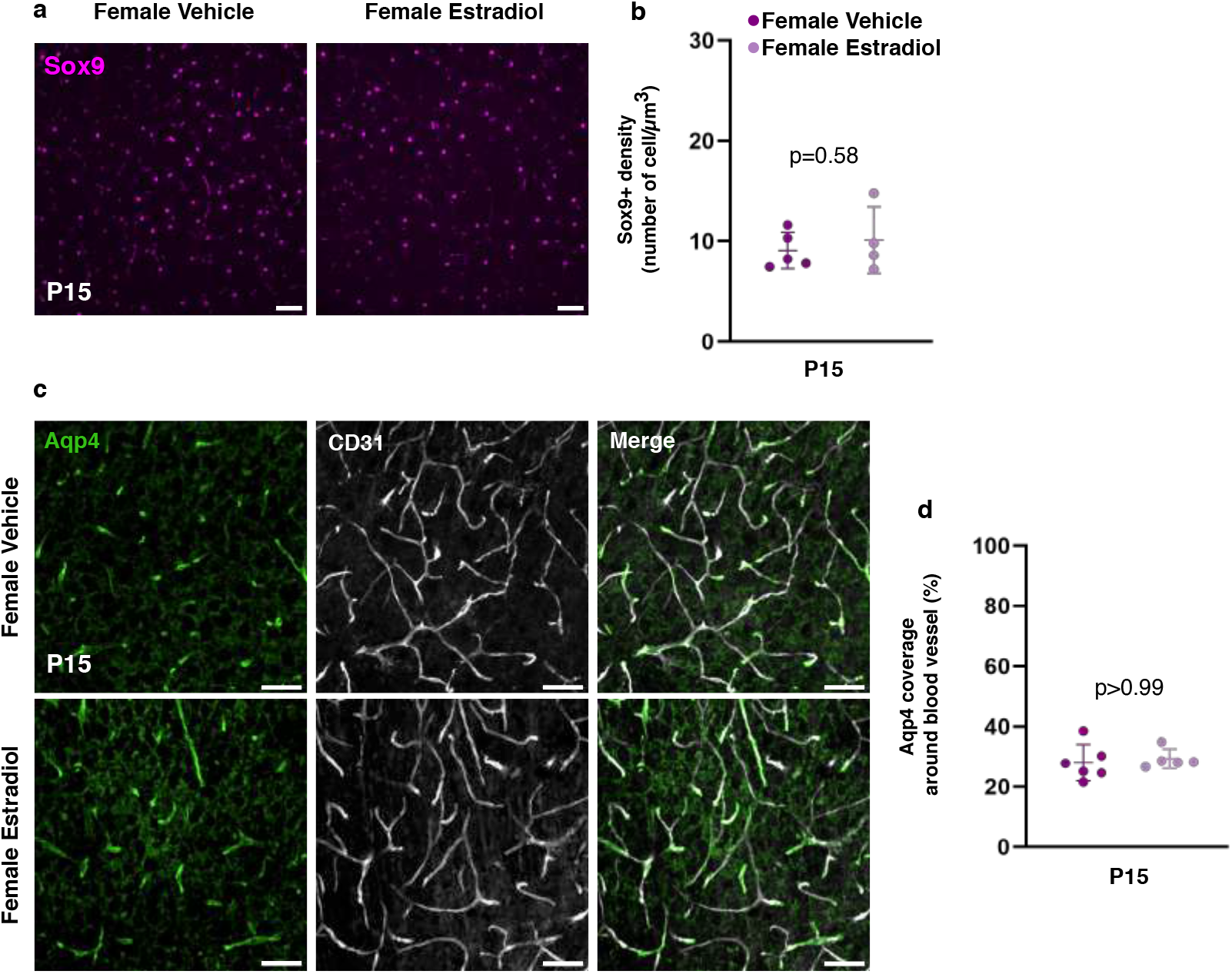
Oestradiol injection on P1 has no effect on astrocyte density or the perivascular level of Aqp4. **a.** Representative images of P15 astrocyte somas immunolabeled for Sox9 (in magenta) in the somatosensory cortex of vehicle- vs. oestradiol-benzoate-treated females. Scale bar: 50 µm. **b.** Quantification of Sox9^+^ cells. n_P15 female-vehicle_ = 5, n_P15 female oestradiol benzoate_ = 4. The data are presented as the mean ± SD and were analysed using a Mann-Whitney test. **c.** Representative spinning-disk images of astrocyte perivascular processes immunolabelled for Aqp4 (in green) in male and female somatosensory cortex sections on P15. Vessels are immunolabelled for CD31 (in grey). Scale bars: 50 µm. **d.** Quantification of perivascular Aqp4 on P5, P15, and P30. n_P15 female-vehicle_ = 6, n_P15 female oestradiol benzoate_ = 5. The data are presented as the mean ± SD and were analysed using a chi^2^ test.

**Supplementary Figure 7:**
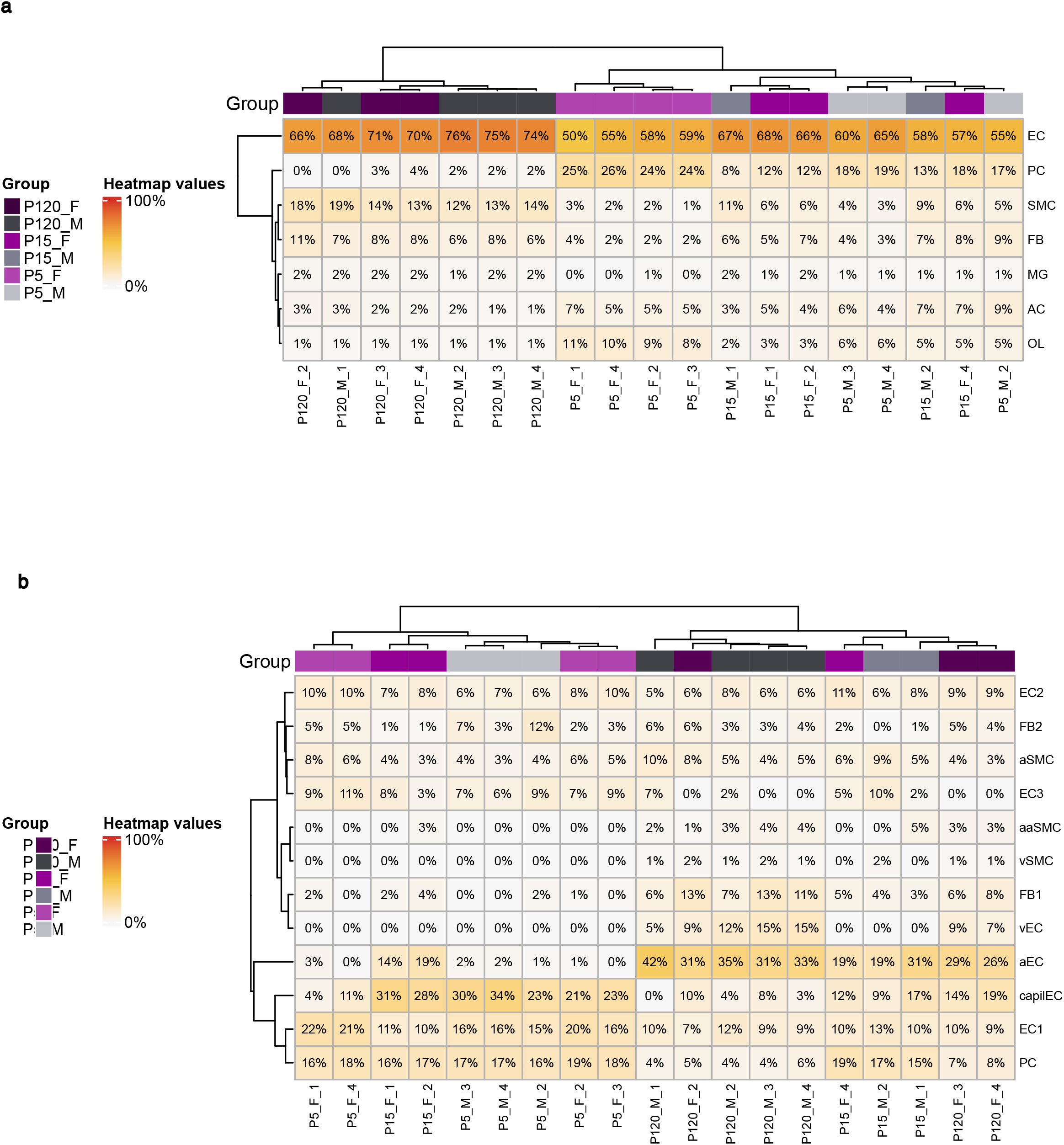
Hierarchical clustering of cell origin of mRNAs sequenced from cortical male and female MVs on P5, P15 and P120. **a.** Vascular and glial cells **b.** Vascular cell subtypes. Cell population signatures were defined using single-cell RNA-Seq data from GEO datasets GSE99058 and GSE98816 ^124,125^.

**Table S1: GVU transcriptome of male and female mice on P5, P15 and P120.** Columns F to P show the percentage of cells expressing each gene, according to ^63,64^. EC, endothelial cell; aEC, arteriolar endothelial cell; capill, capillary endothelial cell; MΦ, macrophage; vSMC, venous smooth muscle cell; aaSMC, arterial smooth muscle cell. Genes are ***expressed*** (TRUE, otherwise FALSE) if they are detected in more than half of the samples and have a fragments per kilobase of transcript per million mapped reads value exceeding that of 98% of the intergenic regions. Genes are considered to have ***enough uniquely mapped reads*** (TRUE, otherwise FALSE) if they have at least 50% of reads mapped on the coding sequence. Genes are considered to be ***regulated*** (value = 1, otherwise 0) if they meet the above criteria (both TRUE) with a false discovery rate ≤ 0.05 and |log2fold-change| ≥ 1.

**Table S2: Clusters corresponding to distinct developmental profiles in male and female GVU between P5, P15 and P120.** M, male; F, female. Clusters MC0 and FC4: synaptic organisation; clusters MC1 and FC5: cell cycle; clusters MC2 and FC3: neuron differentiation; clusters MC4 and FC0: immune response; clusters MC3 and FC2: cytoskeleton organization; clusters MC5 and FC1: detoxification

**Table S3: Genes regulated in opposite directions in males vs. females.**

**Table S4: Reagent list**

**Table S5: Raw data and statistical analyses**

